# Estimating Multiple Latencies in the Auditory System from Auditory Steady-State Responses on a Single EEG Channel

**DOI:** 10.1101/2020.09.27.315614

**Authors:** Lei Wang, Elisabeth Noordanus, A. John van Opstal

**Affiliations:** Department of Biophysics, Radboud University, Nijmegen, 6525 AJ, The Netherlands; Donders Centre for Neuroscience, Radboud University, Nijmegen, 6525 AJ, The Netherlands

**Keywords:** Auditory system, EEG, ASSRs, Apparent latency from phase coherence (ALPC), Multi-cosine stimuli

## Abstract

The latency of the auditory steady-state response (ASSR) may provide valuable information regarding the integrity of the auditory system, as it could potentially reveal the presence of multiple intracerebral sources. To estimate multiple latencies from high-order ASSRs, we propose a novel two-stage procedure that consists of a nonparametric estimation method, called apparent latency from phase coherence (ALPC), followed by a heuristic sequential forward selection algorithm (SFS). Compared with existing methods, ALPC-SFS requires few prior assumptions, and is straightforward to implement for higher-order nonlinear responses to multi-cosine sound complexes with their initial phases set to zero. It systematically evaluates the nonlinear components of the ASSRs by estimating multiple latencies, automatically identifies involved ASSR components, and reports a latency consistency index (LCI). To verify the proposed method, we performed simulations for several scenarios: two nonlinear subsystems with different or overlapping outputs. We compared the results from our method with predictions from existing, parametric methods. We also recorded the EEG from ten normal-hearing adults by bilaterally presenting superimposed tones with four frequencies that evoke a unique set of ASSRs. From these ASSRs, two major latencies were found to be stable across subjects on repeated measurement days. The two latencies are dominated by low-frequency (LF) (near 40 Hz, at around 41-52 ms) and high-frequency (HF) (>80 Hz, at around 21-27 ms) ASSR components. The frontal-central (FC) brain region showed longer latencies on LF components, but shorter latencies on HF components, when compared with temporal-lobe regions. In conclusion, the proposed nonparametric ALPC-SFS method, applied to zero-phase, multi-cosine sound complexes is more suitable for evaluating embedded nonlinear systems underlying ASSRs than existing methods. It may therefore be a promising objective measure for hearing performance and auditory cortex (dys)function. The Matlab scripts for the ALPC-SFS method is available here: https://github.com/ieeeWang/ALPC-SFS-method-Matlab-scripts.

## I. Introduction

Auditory steady-state responses (ASSRs) are stable brain oscillations that are locked to the frequencies present in the periodic envelope of acoustic stimuli ([1] [2]). Studies have indicated that ASSRs may be generated at different levels within the auditory pathway, ranging from as early as the cochlear nerve and subcortical sources ([3], [4]), to the neocortex ([5], [6], [7], [8]). Because of their reproducibility, and involuntary nature, ASSRs have been considered a valid objective biomarker for auditory system disorders that feature abnormal sound processing, as well as for evaluating primary and non-primary auditory cortex function ([9], [10]). Its presence could relate to basic bottom-up sound processing ([11], [12]), as well as to more complex cognitive skills, such as auditory learning ([13]), selective attention ([14],) speech and music perception ([7]), mental disorders ([15]), or illnesses such as tinnitus ([16], [9]). Furthermore, since ASSRs may contain signatures of the neural processing from auditory periphery to cortex, the electrically evoked ASSR from a cochlear implant (CI) could provide an objective measure for the responsiveness of different regions of the auditory pathway of hearing-impaired CI-users ([17]).

It should be noted that ASSRs are due to nonlinear mechanisms in the auditory system (see [18], for a review). At the earliest stages in the auditory pathway, the nonlinear character of neural sound encoding has been evaluated from auditory nerve recordings. A stimulus consisting of multiple pure tones (the carrier frequencies) was shown to result in an envelope spectrum that could be characterized by a unique series of 2^*nd*^-order nonlinear difference frequencies (or ‘beats’) between any pair of carrier frequencies, which all showed up in the auditory nerve response. This indicates that these 2^*nd*^-order nonlinear distortions already arise at, or before, the auditory nerve ([19]). In the ascending auditory pathway, such harmonic complexes might also evoke higher-order nonlinear components, which in turn could give rise to additional ASSRs. In particular, when two pure tones with frequencies *f*_1_ and *f*_2_, are passed through a nonlinear system with system order *R*, a series of combination tones is produced at the output, characterized by *nf*_1_ ± *mf*_2_ (> 0), with *n* and *m* positive integers, such that *n* + *m* ≤ *R* ([16], [20]). So far, few studies have systematically evaluated such higher-order nonlinearities in the EEG responses of the human auditory system.

| Symbol & term | Representation |
| --- | --- |
| $\alpha_{\Sigma_i}$ | current phase angle of output frequencies $\in [0, 2\pi]$ |
| $\varphi_{\Sigma_i}$ | initial phase angle (at $t=0$) of output frequencies $\in [0, 2\pi]$ |
| $\varphi_i^{in}$ | initial phase angle (at $t=0$) of input frequencies $\in [0, 2\pi]$ |
| $\omega_{\Sigma_i}$ | output angular frequencies |
| $\omega_i^{in}$ | input angular frequencies |
| $\tau_e$ | estimate of real latency |
| $\tau_p$ | pseudo latency |
| $\Delta_t$ | the onset delay of a target EEG epoch |
| $C_\tau^i$ | latency consistency index (LCI) of $i$-th ASSR |
| $C_\tau^{sig}$ | 95% confidence limit of LCI |
| $d_\phi$ | the slope of phase lags vs frequencies |
| $\xi$ | relative amplitude gain ($\xi > 0$) |
| $\theta$ | threshold for PLV |
| $\phi(f_i)$ | compute current phase of a frequency by FT |
| $\text{mod}(\cdot)$ | modulus function: remainder after division |
| PLV | phase lock value |
| MPE | mean phase error at the estimated latency $\in [0, 2]$ |
| MSPC | multi-spectral phase coherence |
| ALPC | apparent latency from phase coherence |
| SFS | sequential forward selection |
| MB | monaural beat |

Apart from these nonlinear distortion components, also the latency at which they occur in the EEG provides information of the underlying neural mechanisms, and could reveal the contribution from different intracerebral sources ([21], [6], [7]). Although the auditory system consists of both ascending and descending pathways ([5]), the ASSRs recorded at the scalp are assumed to be mainly due to the ascending system. Therefore, the longer the latency, the higher in the auditory system its generators ([1]).

A steady-state response is characterized by its amplitude and phase. The phase (with certain ambiguity) can be used to extract a so-called ‘apparent latency’ (the slope of the phase versus frequency plot) introduced by Regan (1966). In auditory physiology, the apparent latency is described as ‘group delay’ ([22]) and has been used to measure the latency of the motion of the basilar membrane, the cochlear microphonic, otoacoustic emissions, and the discharge of auditory nerve fibers, although the actual relation of apparent latency to physiological delay is not clear ([1]). Specifically, apparent latency, *τ*, is derived from the relation between the response phase, Δ*ϕ*, and frequency: Δ*ϕ* = 2*πf* · *τ*), and is used as an indirect method to estimate the neural response latency for steady-state stimuli, when a direct measurement is not feasible ([22]). This method, however, suffers from the inherent 2*π* phase ambiguity from the so-called wrap-around effect ([23]), and is only suitable for analyzing low-order ASSR components (e.g., envelope frequencies) when the initial phases are known (so that phase lags can be computed). The more recently-developed multi-spectral phase-coherence (MSPC) method ([20]; or similarly, the cross-spectral coherence (CSC) method, [24]) resolves the phase-ambiguity problem, and allows evaluation of higher-order nonlinear response components. However, the MSPC method relies on prior assumptions regarding the underlying nonlinear systems: knowledge about the system orders is required for evaluating the expected nonlinear interactions, from which response latencies can be estimated. Therefore, the MSPC method is difficult to implement when the number of underlying systems and their orders (and thus, their nonlinear interactions) are unknown. Moreover, when using multiple carrier frequencies, there is an explosion of the number of potential nonlinear interactions: for given nonlinear systems and multi-tone sinusoidal inputs, a recursive method ([25]) can yield the series of output frequencies. The number of output frequencies is a steeply increasing function of the system order *R* and the number of input frequencies, *I*, and its accurate computation is non-trivial because of potentially overlapping distortion products ([26]; see also Appendix C in Supplemental Material).

Thus, the current state-of-the-art methods are limited in estimating multiple latencies for different nonlinear systems from the ASSRs. However, as indicated above, several studies have suggested that ASSR sources can be located at multiple cortical and subcortical regions ([6], [3]), which are likely associated with different latencies. For example, the two-component (brainstem-cortex) model suggests that a cortical generator will dominate low-frequency distortion products (around 45 Hz), whereas a brainstem generator would dominate the higher modulation frequencies (around 90 Hz) ([2], [27], [28]). As the measured EEG will typically contain the components from these multiple generators, a more general hybrid nonlinear model would read:

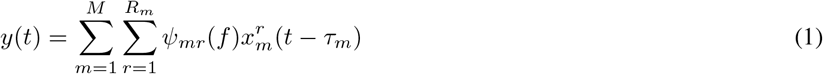

in which the unknown parameters are the number of subsystems *M*, the subsystem order *R_m_*, the subsystem latencies *τ_m_*, as well as the potentially frequency-dependent gains, *ψ_mr_*. Note that, in general, it will not be possible to uniquely identify each nonlinear system, when the underlying nonlinearities and their inputs are unknown. Yet, for certain cases, system-specific information can be obtained from the ASSRs. Figure 1 illustrates a potential simple scenario of such nonlinear models. The EEG is considered to reflect the (linear) sum of the outputs from multiple nonlinear subsystems (here taken, for simplicity, as homogeneous nonlinearities of different orders, *R_m_*, and each at a different latency, *τ_m_*). The output distortion frequencies of these subsystems will differ when they receive non-overlapping inputs (e.g., systems *S*_1_ and *S*_2_). In this case, the latencies of both subsystems can be accurately estimated. In contrast, the output distortion products could be largely overlapping when the systems receive identical inputs, as in *S*_2_ and *S_M_*. In such a case, it will be much harder to estimate the underlying latencies. In this paper, we simulated both cases.

**Fig. 1.**
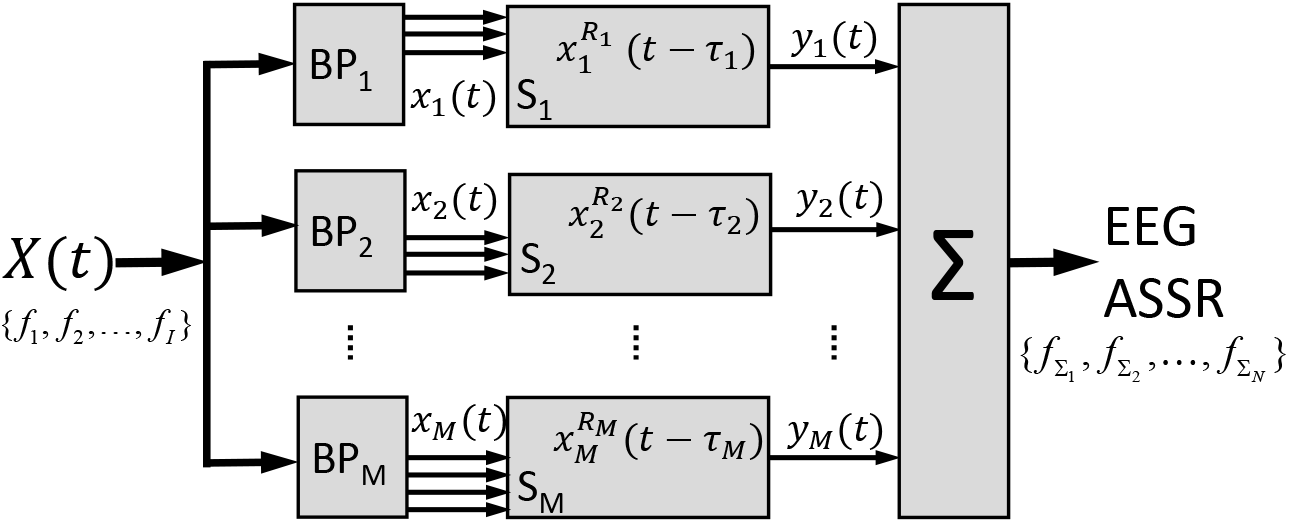
Simplified scenario of M nonlinear subsystems that underlie the measured EEG response. The input signal *X*(*t*) contains *I* frequencies. *BP_m_* represent different band-pass filters, so that the subsystems (with order *R_m_* and latency *τ_m_*) may be driven by the same or by different input frequencies. The EEG contains the linear sum of all subsystem outputs *y_m_*(*t*), with *N* nonlinear distortion components *f*_∑*n*_ (ASSRs).

There are currently no suitable methods to deal with such models, and the the commonly used apparent latency (or group delay) method cannot be readily applied. The apparent latency method requires computation of phase lags for a group of nearby frequencies, which is difficult for higher-order ASSRs, due to (1) the 2*π* wrapping problem (i.e., how many cycles have passed?), (2) the nonlinear distortion of phases for higher-order ASSRs (which could induce a bias within 2*π*, see Appendix C in Supplemental Material), and (3) higher-order ASSRs often cover a large frequency range. Thus, the assumption that a group of target frequencies share a common latency does not apply anymore. As described earlier, the SPC method may partly account for the first two problems through a parametric approach, which then hinges on the assumption of the underlying nonlinear system orders (and their corresponding nonlinear interactions). It then estimates a latency that follows the apparent latency rule (i.e., *τ* = Δ*ϕ*/2*πf*). Here, we applied the MSPC method as an extension of the classical group delay method for higher-order ASSRs. However, the method assumes a common latency for all nonlinear response components with the same system order (e.g., all 2nd order responses), which might not be suitable for the ASSR analysis when the EEG signals arise from multiple subsystems (as illustrated in Fig. 1).

We here propose and test an alternative, nonparametric, semi-heuristic method that requires few prior assumptions regarding the underlying EEG sources, and aims to objectively estimate multiple potential latencies from the ASSRs. We named our method ‘apparent latency from phase coherence’ (ALPC-SFS), which is applied to a specifically designed stimulus complex, and implements a heuristic sequential forward selection (SFS) algorithm ([29]) to identify the components of each subsystem. Our method can estimate multiple latencies corresponding to more than one underlying source by using only one EEG experiment. Our ALPC-SFS method does not have to make prior assumptions about the number of underlying systems, their input frequencies, or their system orders. To test our method, we performed simulations for several scenarios, and recorded EEG responses from ten normal-hearing participants to superimposed tones that evoked high-order ASSRs during passive listening.

We also compared our ALPC-SFS method with the existing MSPC method, for both the simulation data and the EEG recordings. Both methods can be viewed as extensions of the classical group-delay method for higher-order ASSRs, as both follow the apparent latency rule. They will achieve an identical latency estimate if only one common latency exists. However, when multiple potential latencies underlie the EEG, our ALPS-SFS method will outperform the MSPC method (and the group delay method), achieving correct latencies for each subsystem, and identifying the associated frequencies that drive each subsystem (see Fig. 3).

## II. Methods

To estimate multiple latencies from recorded ASSRs, we developed a two-stage procedure that consists of the nonparametric ALPC estimation method, followed by the heuristic SFS algorithm. The ALPC method can apply phase compensation on stimuli with arbitrary initial phases (as used in [20]), or it can use a much simpler time compensation (TC) procedure for a special stimulus complex, in which the initial phases of all stimulus frequencies, and thus their nonlinear distortion products, are set to zero. Note that the former requires prior information regarding the underlying nonlinear orders of systems and the corresponding nonlinear interaction (see Appendix A in Supplemental Material). In contrast, ALPC with TC requires no such information, and thus can be viewed as a nonparametric method (see the right-hand side of Fig. 3). We combined ALPC with SFS and TC to identify multiple latencies from the ASSRs.

### A. Problem description

The EEG is considered as a time-series signal, *S*, containing a number of distortion frequency components {*f*_∑1_, *f*_∑2_,..., *f*_∑*N*_ with corresponding angular frequencies *ω*_∑*i*_ = 2*πf*_∑*i*_, which are the outputs of a nonlinear system. Assuming that *N* frequency components of them have a common, unknown, delay *τ* relative to their initial phase *φ*_∑*i*_ at t=0 (potentially having a common generator), we compute the current phase angle *α_∑i_* of each component by taking the Fourier transform (FT), where *α*_∑*i*_, φ_∑*i*_ ∈ [0, 2*π*]. They follow:

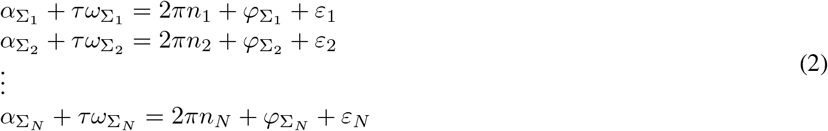

where *n_i_* are unknown integers, i.e., the number of cycles that a component is delayed, and *ε_i_* is the phase error (PE), caused by random noise.

To estimate the common latency, we minimize the mean-squared phase error across *N* frequencies ([30]):

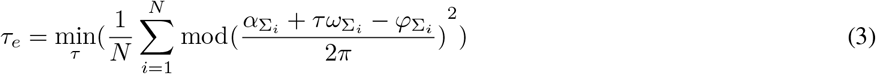

where mod(·) is the modulus between 0 and 2*π*. If the nonlinear system order, *R* ≤ 2, the initial phases are straightforward to measure from the input stimulus by extracting the phases from the envelope of the input signal. However, when the systems order is unknown, and higher than three, this method will fail, because the unknown *φ*_∑*i*_ are endowed with the wrap-around problem (i.e., unknown *n_i_*).

### B. General framework of ALPC

First, to avoid that a discontinuous ‘jump’ (from 0 to 2*π*) would affect the averaging in (3), we represent the components of (2) by Euler’s formula according to: 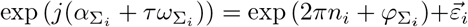 or, equivalently, 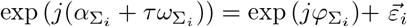, where 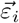 is a vectorized phase error (PE) and its length 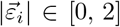. We then minimize the cost, defined by the mean length of PE (MPE) across N frequencies, to estimate *τ_e_*:

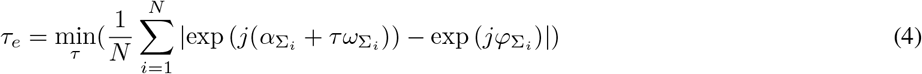

where | · | is the length of a complex vector. Using the estimated *τ_e_*, we can compute *n_i_* and the absolute phase lag of each frequency by 2*πn_i_* + *φ*_∑*i*_ – *α*_∑*i*_ (equal to *τ_e_ω*_∑*i*_ − *ε_i_*). Plotting the absolute phase lags for each frequency component as a function of frequency should yield a line with a slope *d_ϕ_*, where the relationship between *τ_e_* and *d_ϕ_* follows the apparent latency rule ([1]):

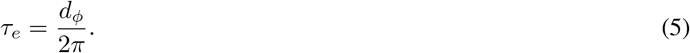

In (4), we estimate *τ_e_* by searching within a period *T* of signal *S*, where *T* is the smallest common multiple of the periods of frequency components {*f*_1_, *f*_2_,..., *f_N_*}. Then, the solutions for the latency are given by a set {*τ*|*τ* = *τ_e_* + *mT*}, with *m* an arbitrary integer. In a particular application, a *prior* for the possible range of latency values can be imposed to constrain a unique value for *τ_e_*.

#### 1) Time compensation (TC) for stimuli with fixed initial phases

To avoid the need for computing the unknown *φ*_∑*i*_ when using stimuli with arbitrary initial phases (which constitutes ‘phase compensation’, see Appendix A in Supplemental Material), we here propose to use a more practical stimulus complex: a multi-cosine stimulus, in which all initial phases are set to zero at t=0. With such a stimulus complex it is straightforward to apply time compensation (TC) for an arbitrary ongoing interval (starting at t>0) in the following way (see Fig. 2).

**Fig. 2.**
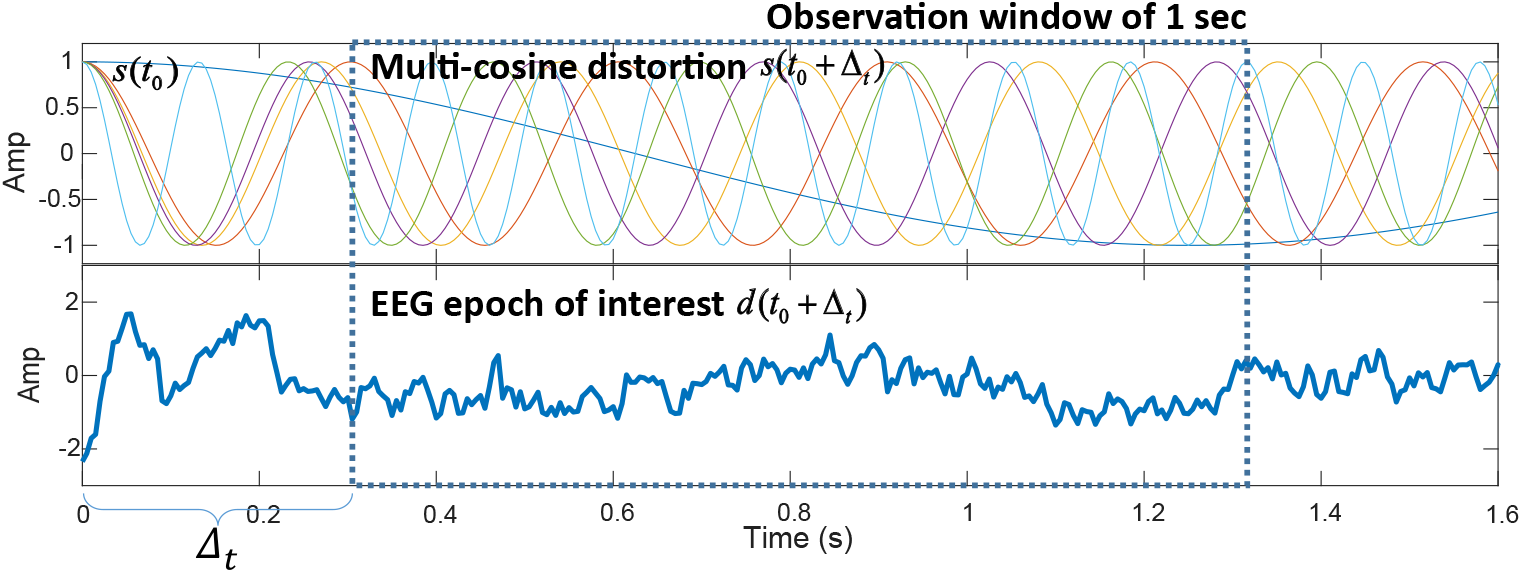
Illustration of time compensation (TC) on a single-channel EEG. The stimulus complex consisted of a superposition of four cosine pure tones, at 461, 500, 504 and 537 Hz, starting at t=0 with initial phase 0. Here, the six distortion frequencies (i.e., the 2^nd^ order system products) in the top panel at [4, 33, 37, 39, 43, 76] Hz, are shown and time-scaled by a factor 0.1, for illustrative purposes. Note that all six distortions also start with zero phase at t=0 (because of the good property of cosine; see Appendix C in Supplemental Material). The EEG response to the stimulus complex is shown in the bottom panel (1^st^ trial on EEG electrode P8, subj#1) and has been down-sampled to 200 Hz for easy visualization. The first Δ_t_ = 0.3 s of the EEG signal is excluded from analysis because of the initial non-stationary ERP components (i.e., non-ASSR).

For a given multi-cosine input stimulus, 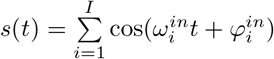, with all initial phases 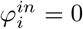 at *t*_0_ = 0, the ongoing stimulus interval, starting at *t*_0_ + Δ_*t*_, is denoted as *s*(*t*_0_ + Δ_*t*_). The delayed EEG signal (the ASSR) *d*(*t*) is considered to be the output of an unknown nonlinear system with an unknown latency, *τ_e_*, i.e., *d*(*t*) = *F*(*s*(*t* – *τ_e_*)), where *F*(·) represents the unknown nonlinear mapping. The EEG epoch of interest will hence follow:

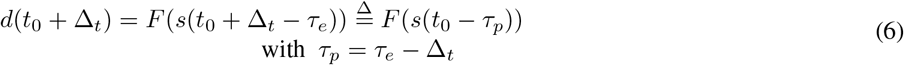

Therefore, from the ASSR we can estimate the pseudo-latency *τ_p_* between *d*(*t*_0_ + Δ_*t*_) and *s*(*t*_0_). Since 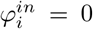, the corresponding initial phases *φ*_∑*i*_ of all high-order responses are also zero (i.e., 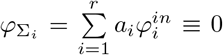, see Appendix A and C in Supplemental Material). Moreover, it can be shown that for zero-phase sine inputs, all even-order difference interactions (e.g., *f_n_* – *f_m_*, and 2*f_n_* – 2*f_m_*) will yield the same initial phase of *π*/2 rad, so that the method can also be applied to these components (after changing the initial phase to *π*/2; see Appendix C). Therefore, from (4) and (6), we can estimate the pseudo latency *τ*p. Finally, we apply TC to obtain the actual latency:

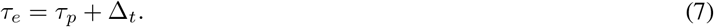

In our study, Δ_t_= 0.3 s, in order to exclude the non-stationary event-related potentials (ERP) in the EEG after stimulus onset (see Fig. 2). The device jitter (resulting to be < 0.5 ms) involved a small trigger jitter of the EEG recording system, and a fixed travel time of the acoustic signal through the silicon tube to the ears. Thus, the estimated latency had around 1-ms system error, which was deemed acceptable for the estimated ASSR latencies.

#### 2) Latency consistency index (LCI)

Considering that the estimated latency may vary due to additive noise on the nonstationary signals, we quantified its consistency over *K* time epochs. We defined the latency consistency index (LCI) for the *i^th^* output frequency *f*_∑*i*_ as 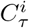.

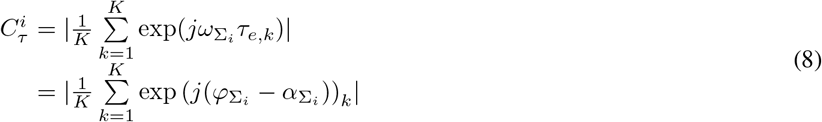

where *τ_e,k_* is the estimated latency for frequency component *f*_∑*i*_ on the *k^th^* epoch, and it follows *ω*_∑*i*_ *τ_e,k_* = 2*πn_i_*, + *φ*_∑*i*_ – *α*_∑*i*_ from (2), assuming Gaussian noise with zero mean. The LCI is hence mathematically equivalent to the phase-coupling strength used in MSPC ([20]), which shows that the higher the phase-coupling strength between inputs and outputs, the more stable the estimated latency. We selected the duration of each time epoch (without overlap) as multiples of the periods of the ASSR frequencies (e.g., integer seconds, given integer ASSR frequencies) so that the initial phase *φ*_∑*i*_ remains the same across epochs. In this case, *φ*_∑*i*_ ≡ 0 when the cosine carriers have their initial phases set to zero. Thus, the LCI no longer relies on the stimulus frequencies, and reads:

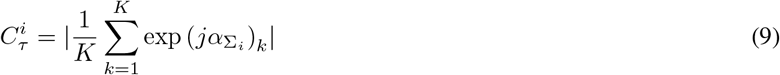

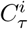 can vary between 0 and 1. If the latency is perfectly stable over time epochs, 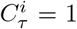. Otherwise, if no stable latency exists, 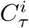 is statistically indistinguishable from zero. Therefore, LCI can be used to determine if an ASSR component can be used to estimate latencies. The theoretical threshold for significance is 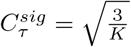 ([31]).

The null hypothesis is that the phase of an ASSR component is randomly distributed in the interval [0, 2*π*]. If 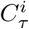 exceeds 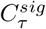, the null hypothesis is rejected, and the ASSR component is considered to be significant. 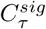 approximates the 95% confidence limit, as verified by Monte Carlo simulation ([20]). In our analysis, only significant ASSR components were considered for estimating latencies.

#### C. ALPC with sequential forward selection (SFS)

A particular set of ASSR components may have a common latency that lead to a low cost of the MPE in (4). In addition, the set of ASSR components may contain several subsets that correspond to different latencies, which would be indicative for different underlying sources. To automatically identify a frequency subset from all significant ASSR components that belong to a common latency, we applied the heuristic SFS algorithm. SFS is commonly used as an alternative to a greedy algorithm, to reduce computational complexity. SFS starts with a frequency (or frequency pair), and then sequentially adds another frequency (from the remaining set) that results in the smallest increase of the cost (i.e., MPE in a range of [0 - 2]). The details of the SFS algorithm are described in ([29]) and ([32]).

The termination rule of SFS should be carefully chosen, depending on the data at hand. Here, we used the following procedure. When starting from a single frequency, SFS continued as long as the following two criteria were met: (i) the increase of the MPE of the current step remained < 0.1, (ii) the current MPE < 0.5. Otherwise, SFS terminates. If SFS started from a pair of frequencies, it continued when the following three criteria were met: (i) the difference of estimated latencies between the current and the previous step <5 ms, (ii) the step increase of MPE < 0.1, (iii) the current MPE < 0.5. Otherwise, SFS terminates. These criteria may be further tuned for the particular EEG data set.

Figure 3 summarizes the pipeline for estimating the apparent latency from the EEG by the MSPC and ALPC-SFS methods, respectively. Depending on the initial phases of the carriers (either arbitrary, or all zero), one can use either a parametric (the left-hand side, MSPC, or group delay) or a non-parametric (the right-hand side; ALPC-SFS) approach to estimate the underlying latency. For the parametric approach, phase compensation (see Appendix A in Supplemental Material) can be used to estimate the unknown initial phases of the ASSRs, which are fed to the MSPC algorithm to estimate the dominant latency. Note that the phase compensation method requires prior assumptions regarding the nonlinear system. For the non-parametric approach as proposed in the present paper, ALPC with SFS can estimate multiple potential latencies, by selecting the involved ASSR frequencies for each subsystem.

**Fig. 3.**
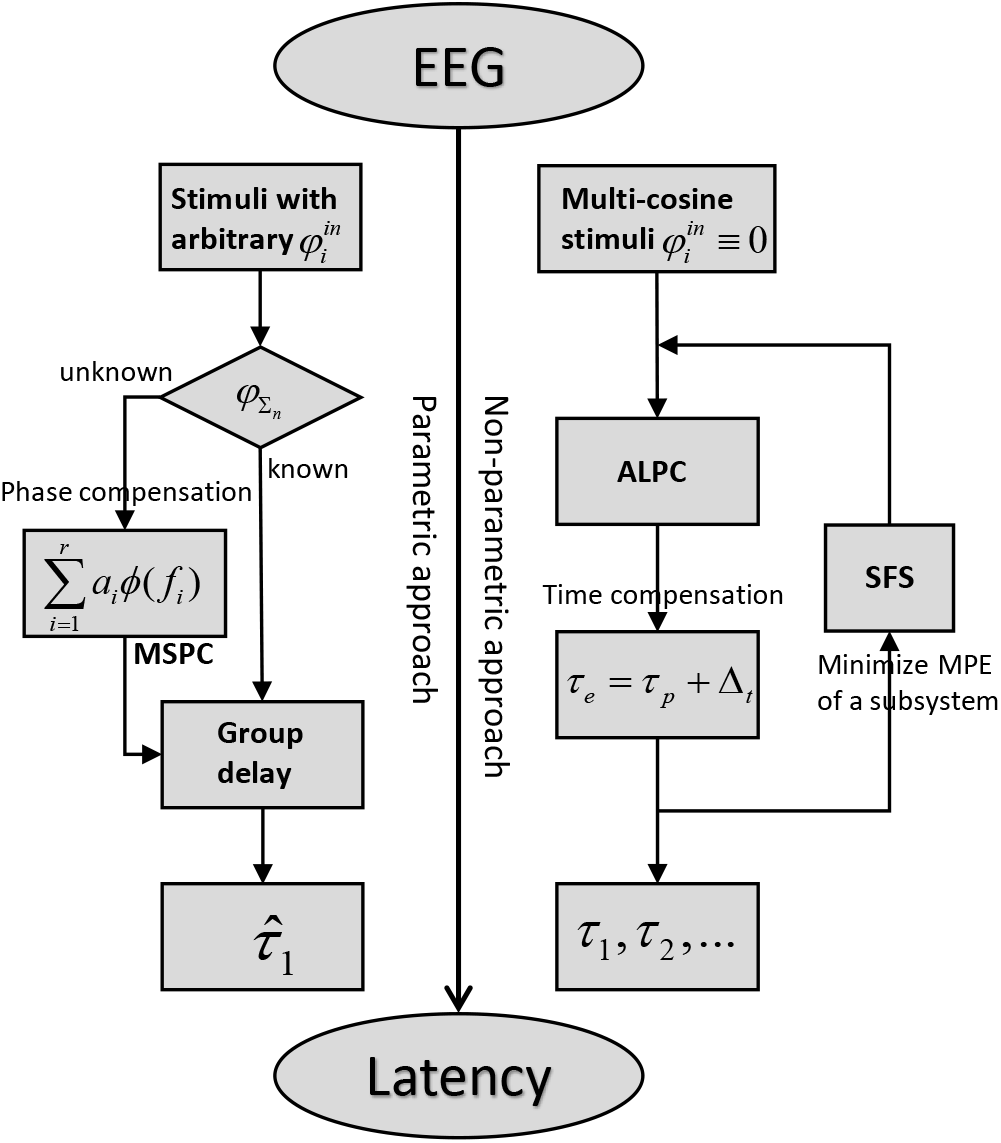
Pipeline for estimating the apparent latency from the EEG signal, using the parametric MSPC (left) or non-parametric ALPC-SFS (right) approach. 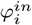 represents the initial phase of each frequency component in the stimuli (input), and *φ*∑_*n*_ represents the initial phase of each ASSR frequency component (output). By using multi-cosine stimuli with their initial phases set to zero, the ALPC-SFS method (right) uses time compensation by Δ_t_ to avoid the more complex phase-compensation procedure of the MSPC method (left; see Appendix A in Supplemental Material), which computes phase lags of the ASSRs. Moreover, the former can identify potentially multiple latencies, [*τ*_1_, *τ*_2_, ⋯], instead of only one (biased) dominant latency, 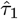.

## III. EEG experiments

### A. Experimental design and stimuli

We performed EEG recording experiments on 10 normal-hearing adults (1 female, age 29 ± 13 y) (hearing thresholds <20 dB HL). Each subject was measured twice, using the same stimuli on different days (i.e., >48 hours interval). The stimulus at the left ear was a superposition of two tones of 461 Hz and 500 Hz at 81 dB SPL, and two tones of 504 Hz and 537 Hz at 56 dB SPL. The stimuli at the right ear consisted of the reverse pattern: two loud tones of 504 Hz and 537 Hz at 81 dB SPL, and two weaker tones of 461 Hz and 500 Hz at 56 dB SPL. The stimuli were fed into both ears simultaneously. The four frequencies were chosen such that the difference frequencies and higher-order distortions were all unique (see Fig. A2 in the Supplemental Material).

In the EEG experiments described here, all input tones were sines with their initial phases at t=0 set to zero. In Appendix C of Supplemental Material, we demonstrate that such carriers will yield irregular phase-shifts for all higher-order distortion products, except for the even-order difference interactions (e.g., *f*_1_ – *f*_2_, and 2*f*_1_ – 2*f*_2_). The initial phase of these even-order distortion products will remain the same. Thus, when only even-order difference interactions in the ASSRs are considered (as in this study), APLC-SFS with TC can also be applied for zero-phase sine stimuli.

In each trial, stimuli with a duration of 12.3 s were played after a silent break with a random duration between 2 and 3 sec. It has been reported that the primary auditory cortex responds within 20 ms to stimulus changes and integrates stimulus features over a period of about 200 ms, which is the pre-steady-state phasic response phase ([33]). Therefore, the first 300 ms of the EEG of each trial was excluded from analysis to avoid the contributions from phasic event-related potentials (i.e., non-ASSRs). We expect the ASSR latencies to remain constant during the subsequent 12 s of the ASSR stage, which will allow for an accurate estimate of the neighboring SNR (see section E). We repeated 100 trials while recording the scalp EEG with a 64-channel cap with Ag/AgCl electrodes, with the listener sitting in an anechoic chamber, watching a silent video. The EEG signals were digitized at a sampling rate of 2000 Hz by a Refa amplifier system (TMSi, Twente Medical Systems International B. V., the Netherlands), and stored for offline analysis. At the start of each recording session, the impedances of all EEG electrodes were carefully set to values below 10 *k*Ω to ensure good contacts between electrodes and scalp. During the session, these values were regularly checked, and adjusted when deemed necessary. Stimuli were generated by TDT (Tucker-Davis Technologies, USA) system 3 hardware, and presented through ER3C insert earphones (Etymotic Research, Elk GroveVillage, IL), which were connected to the participants ears via 30 cm-long plastic tubes and foam earplugs.

The experiments were approved by the ethics committee at Radboud University and performed in accordance with the human experiment guidelines and regulations of Radboud university. We confirm that informed consent was obtained from all subjects.

### B. Preprocessing of EEG

We applied a three-stage preprocessing protocol on the raw EEG recordings, after first removing the three EEG electrodes above the eyes (FP1, FPz, and FP2) because they were contaminated by eye-blinks and eye-movement artifacts. First, the raw EEG signals were re-referenced to a common average reference (CAR) to reduce the noise from the grounding electrode ([34]). Compared with re-reference in relation to a certain EEG electrode that may not be ‘neutral’ to ASSRs, the CAR montage will keep the phases of ASSRs unchanged on most EEG electrodes. Second, each EEG channel was filtered with a zero-phase shift filter ([35]) that consisted of a 10th-order Butterworth high-pass filter with a cutoff frequency of 1 Hz, and a notch filter at 50 Hz to remove the line noise. Third, to exclude severely contaminated EEG trials from further analysis (mainly by EMG artifacts, which may cause false detections or decreased SNRs of ASSR components), we developed a method to automatically exclude bad EEG trials. To that end, we computed the mean and the variance of the peak-to-peak amplitude range as indicators on each EEG trial. We excluded EEG trials whose indicators were positive outliers on 100 trials, i.e., samples > Q3+l.5*IQR. (Interquartile range (IQR) = upper quartile (Q3) – lower quartile (QI)). In this way, up to 5% of all trials were automatically excluded from further analysis.

### C. Epoch filtering by phase lock value (PLV)

We tested whether excluding poorly phase-locked epochs would improve the accuracy of the phase estimates, and latencies, for the frequencies of interest. To that end, we divided each trial into twelve one-second epochs, and computed their phase-lock value (PLV ∈ [0,1]) to each frequency of interest ([36]). PLV = 0 represents no phase locking between the frequency band of EEG and the frequency of interest, while PLV = 1 represents the complete phase locking. Epochs with a PLV value below a empirically selected threshold, *θ* (taken as *θ* = 0, 0.4, 0.6, or 0.8) were excluded. Finally, we computed the phase values on the remaining epochs by using two different averaging methods: AVG EEG, and AVG phase (described below).

### D. Phase extraction of EEG

The accuracy of the extracted phases of the frequency components is important for estimating latency. Previous studies often average the EEG signals across a number of trials to reduce the background noise, so that ASSRs have a higher SNR ([37]). Here, we employed two different methods, ‘AVG EEG’ and ‘AVG phase’ to extract the phase of a target frequency. The ‘AVG EEG’ method averages filtered EEG signals of all trials and subsequently computes the phases of the average EEG. The phase values from this method tend to be dominated by epochs with strong ASSRs (i.e., with large amplitudes). Instead, the ‘AVG phase’ method averages the extracted phases from all EEG epochs with a duration that is an integer multiple of the signal period (1 sec in this study), without considering amplitude information (as used in [20]). For this method, all EEG epochs thus contributed with equal weight to the estimate. For comparison, we used both methods to extract the phases on each EEG channel independently.

### E. SNR on target frequencies

It is difficult to separate the signals generated by ASSRs from background noise in the EEG on a single-trial basis. However, after averaging the EEG across a number of trials (100 trials in this study), the power of phase-locked frequencies is enhanced, whereas the power of the background EEG is suppressed due to random phases. To quantify the SNR of each ASSR frequency component, we defined the neighboring SNR as ([3]):

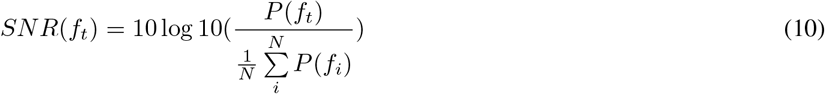

where *P*(*f_t_*) is the power of the target frequency bin *f_t_*. The *f_i_* is a neighboring frequency bin within a small range (±0.5 Hz) of *f_t_*. For a 12 s EEG trial, the width of a frequency bin is 1/12 Hz and *N* =12. Compared with the frequency spectrum of the EEG, the neighboring SNR is robust to the pink noise present in EEG signals (see Fig. 9).

Note that the defined neighboring SNR is not an unbiased estimate of real SNR. In fact, the expected neighboring SNR *E*(*SNR*(*f_t_*)) = *SNR* +1 ([38]). More specifically, the defined neighboring SNR follows a F distribution with (2, 2*N*) degrees of freedom (when the target frequency bin contains no signal), where *N* is the number of neighboring frequency bins. Given a type-I error (e.g., *α* = 0.05) from the F test, the significance threshold θ is a function of the number of neighboring frequency bins ([39]), i.e., *θ* = finv(1 - *α*, 2, 2*N*) by using the MATLAB (2018a) function ‘finv’. For *N* =12, the corresponding thresholds for a significance level p of 0.05 and 0.01 are 3.401 (5.318 dB) and 5.614 (7.492 dB), respectively.

## IV. Results

### A. Simulation results

To illustrate the potential of ALPC with SFS in combination with zero-phase cosine stimuli, we simulated different models and scenarios (cf. Fig. 1). In these examples, two parallel nonlinear sub-systems with different delays could have different, or (partially) shared inputs.

#### 1) Example 1: Two pure second-order subsystems with non-overlapping inputs

Figure 4 shows the simulation results of our ALPC-SFS algorithm for the simple example of Eq. 11. The total input signal, *x*(*t*), consisted of five frequency components: (17, 21, 27, 41, 49) Hz, where *x*_1_=(17, 21, 27) Hz were fed to a pure second-order subsystem *y*_1_, and *x*_2_=(41, 49) Hz to a similar subsystem *y*_2_. In this model, system *y*_1_ generated the distortion products {2*f_i_*, *f_i_*±*f_j_*} = {4, 6,10, 34, 38,42,44,48, 54} Hz, and system *y*_2_ yielded the non-overlapping components {8, 82, 90, 98} Hz. Subsystems *y*_1_ and *y*_2_ had different latencies *τ*_1_ = 51 ms and *τ*_2_ = 21 ms, respectively. *Y* is the combined output signal, (representing the EEG) with additive Gaussian white noise (WGN), *ε*(*t*). We added noise with different variances, such that the SNRs were 5, 0, −5, −10, −15, and −20 (dB) (using MATLAB’s function ‘awgn’).

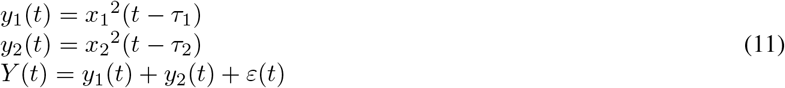

**Fig. 4.**
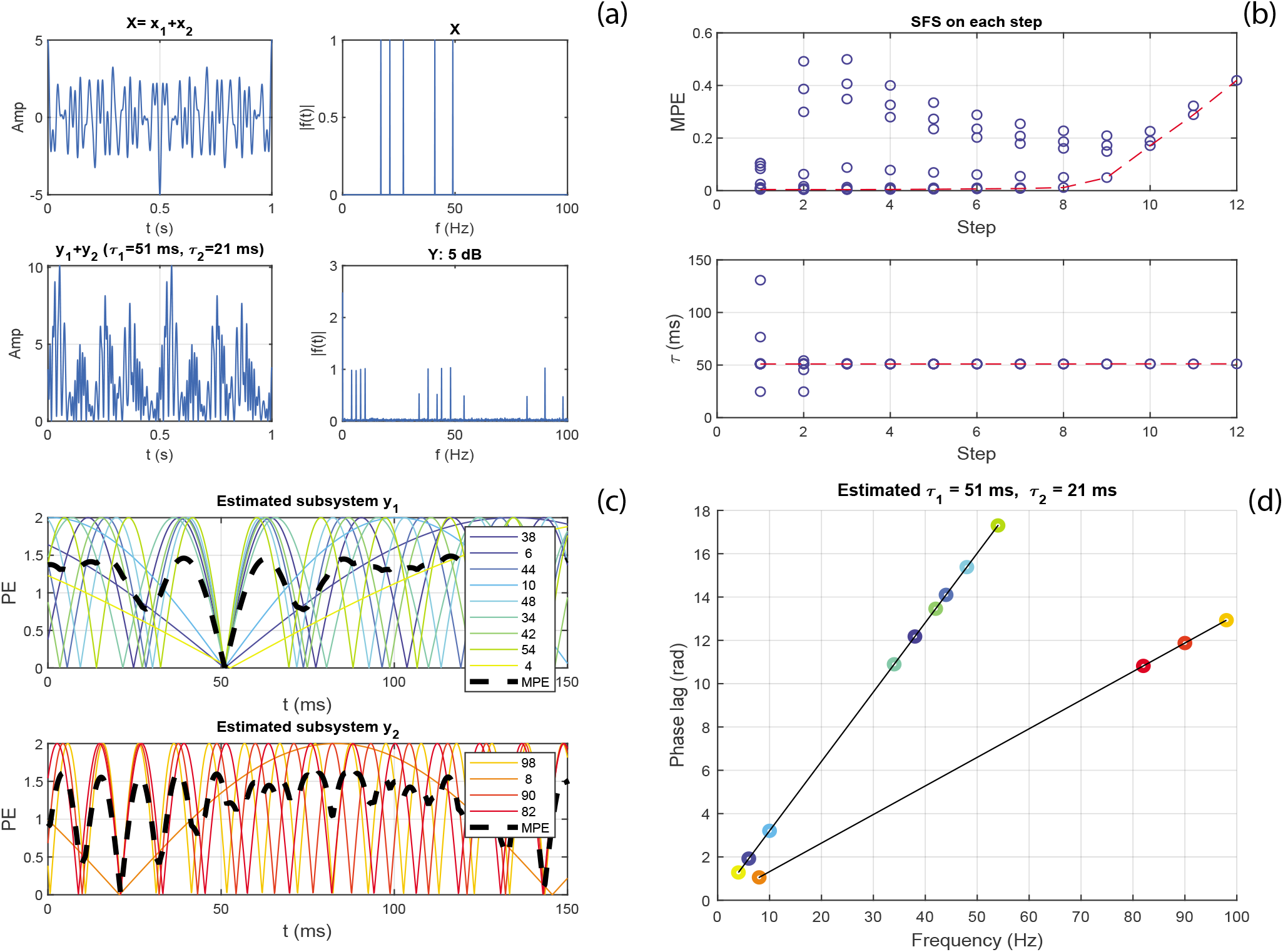
Simulation of the ALPC-SFS method for the system of Eq. 11. The two subsystems were successfully separated from the mixed nonlinear response, and their latencies were estimated correctly. (a) Input signal X (sum of five cosine signals with zero phases) and their mixed total response, *Y*. (b) SFS without applying the termination criterion. Each circle represents a candidate frequency and the dashed line crosses the candidate frequencies with the minimum MPE for each step (top panel) and their corresponding latencies (bottom panel). (c) Phase error (PE, in radians) of each selected frequency over time. The MPE (bold dashed line) reaches its minimum (close to 0) at the estimated latency. (d) Plot of phase lags determined from the estimated latency. The slopes of the best-fit lines correspond to the latencies.

To estimate the unknown latencies, we assume that only the total input (stimulus) and output (EEG) are available, but that the information regarding the subsystems (e.g., input frequencies of each subsystem, and system orders) is unknown. Figure 4 shows that our ALPC-SFS method separated the mixed outputs into the appropriate contributions from the two underlying subsystems, and correctly estimated their latencies. Here, we took for the additional signal delay, Δ*_t_* = 0, for simplicity, and signal Y had an SNR of 5 dB. Figure 4a shows the input and output signals, and their spectrum. Figure 4b illustrates the SFS procedure, starting from a randomly selected frequency component (here: 38 Hz) from the output frequency set (i.e., those frequencies whose power are higher than the background). The upper panels show the mean phase errors (MPEs) of all candidate frequencies on each step in the SFS, the bottom panel shows the corresponding estimated latencies of candidate frequencies at each step. The MPE started to rise significantly after step 8, where a termination criterion of SFS came into action (see Methods). Thus, the first nine (8+1) frequencies were selected, yielding an estimated latency of 51 ms. Subsequently, SFS was performed on the remaining frequency set, which gave an estimated latency for a second subsystem of 21 ms. Without SFS, an estimate based on all frequencies would result in only one dominant latency (51 ms), as shown for step 12 (Fig. 4b).

The relationship between SNR and accuracy of the estimated latencies is provided in Table A1 of Supplemental Material. We also considered a scenario for which Δ*_t_* was not zero. We first estimated the pseudo latencies and applied TC to determine the actual latencies. The pseudo latency uses the property that the initial phases of the nonlinear distortions of zero-phase cosine inputs remain the same (see Appendix C in Supplemental Material for details). In this way, we do not need to apply phase compensation. However, when Δ*_t_* exceeds the actual latency, the corresponding pseudo latency will be negative (Eq. 6), yielding a negative slope for the phase-lag vs. frequency relation (See Fig. 5).

**Fig. 5.**
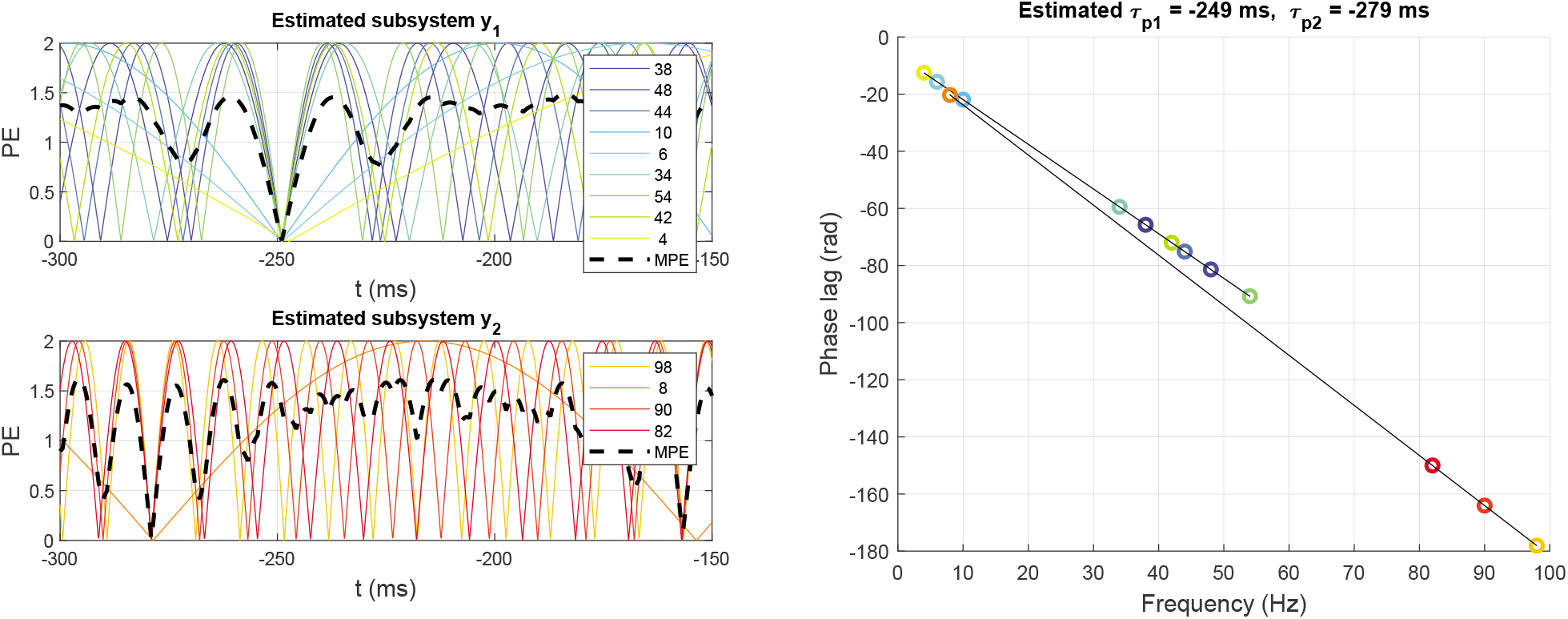
The process of estimating the pseudo latency when the stimuli are delayed (here: Δ_t_ = 300 ms), like in Fig. 2. After applying TC (Eq. 7), the estimated latencies are correctly estimated as 51 and 21 ms, respectively, with its MPE close to zero at these latencies. Notice that the negative slopes in the right-hand panel correspond to the negative pseudo latencies of −249 and −279 ms, respectively (see Methods).

For comparison, we also applied the MSPC method ([20]) to this simple model. Also this method estimates a system latency by minimizing the MPE over a time window. It requires the total input (stimuli) and output (EEG), and has to make an assumption about the underlying system orders. In case of a correct selection of inputs and outputs and system-order information, MSPC may perform equally well as ALPC-SFS (Fig. 3). However, when the sub-systems inputs and outputs are not known a priori, MSPC may fail to correctly identify the systems and make a biased estimate. Figure 6 demonstrates this for the same settings as in Fig. 4. The MSPC method estimated only one biased latency (close to the dominant one) for the assumed 2^*nd*^-order system. However, it failed to find the second hidden subsystem, which produced fewer nonlinear distortion products.

**Fig. 6.**
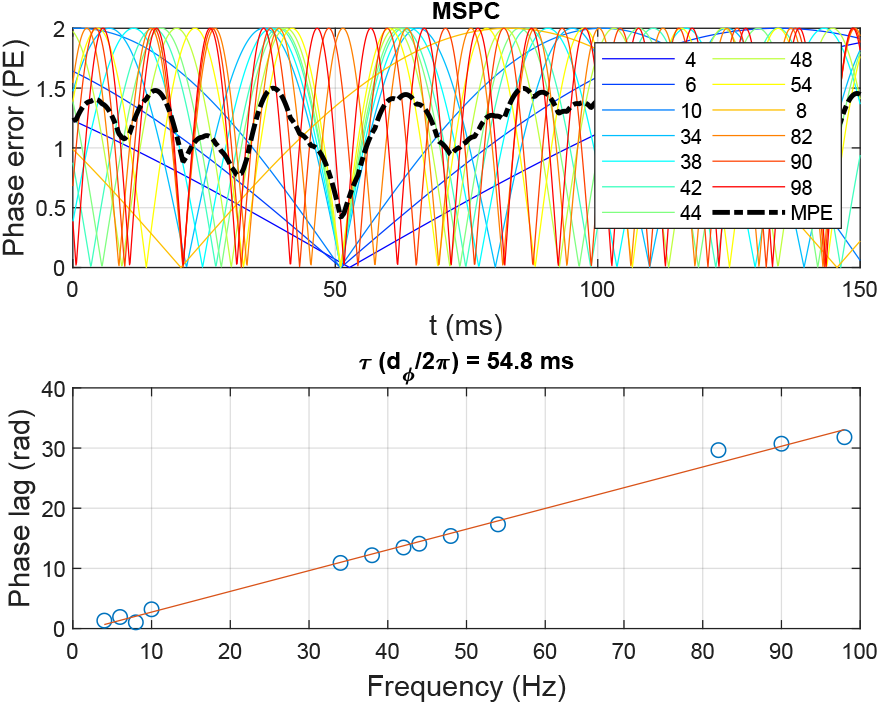
The MSPC method estimates a system’s latency for the same input X and output Y as in Fig. 4. It assumes that the system order equals two. It estimates one biased latency of 54.8 ms with its corresponding MPE of 0.42 (MPE ∈ [0, 2]).

#### 2) Example 2: Models containing multiple system orders

We also applied both methods to a more complex model, in which each subsystem contained a second- and a third-order nonlinearity:

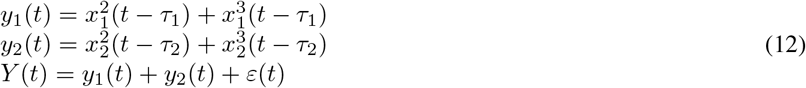

The total input signal *X*(*t*) contained four frequency components: {37, 43, 38, 46} Hz, where *x*_1_=(37, 43) Hz were fed to subsystem *y*_1_, and *x*_2_=(38, 46) Hz to subsystem *y*_2_. In this example, subsystem *y*_1_ generated the following distortion product set: {6, 74, 80, 86, 31, 37, 43, 49, 111, 117, 123, 129} Hz, and subsystem *y*_2_ yielded the non-overlapping set: {8, 76, 84, 92, 30, 38, 46, 54, 114, 122, 130, 138} Hz. For both frequency sets, the first four entries are 2^*nd*^-order distortion products and the remaining components are 3^*rd*^-order outputs. Subsystems *y*_1_ and *y*_2_ had latencies *τ*_1_ = 51 ms and *τ*_2_ = 21 ms, respectively. Y is the combined output signal, representing the EEG.

To estimate the latencies, we assumed that only the total input (*X*) and output (*Y*) were available, and that there was no prior information about the subsystems. Figure 7 shows the estimated latencies for the ALPC-SFS and MSPC methods. In this example, signal Y had an SNR of 5 dB. Without knowing the information of both subsystems *y*_1_ and *y*_2_, ALPC-SFS correctly estimated the latencies and separated the two original subsystems. In contrast, the MSPC method relied on an explicit assumption regarding the order of potential subsystems. However, as in this example, within a given system order, not all distortions are associated with a common latency. As a result, the MSPC method made biased estimates.

**Fig. 7.**
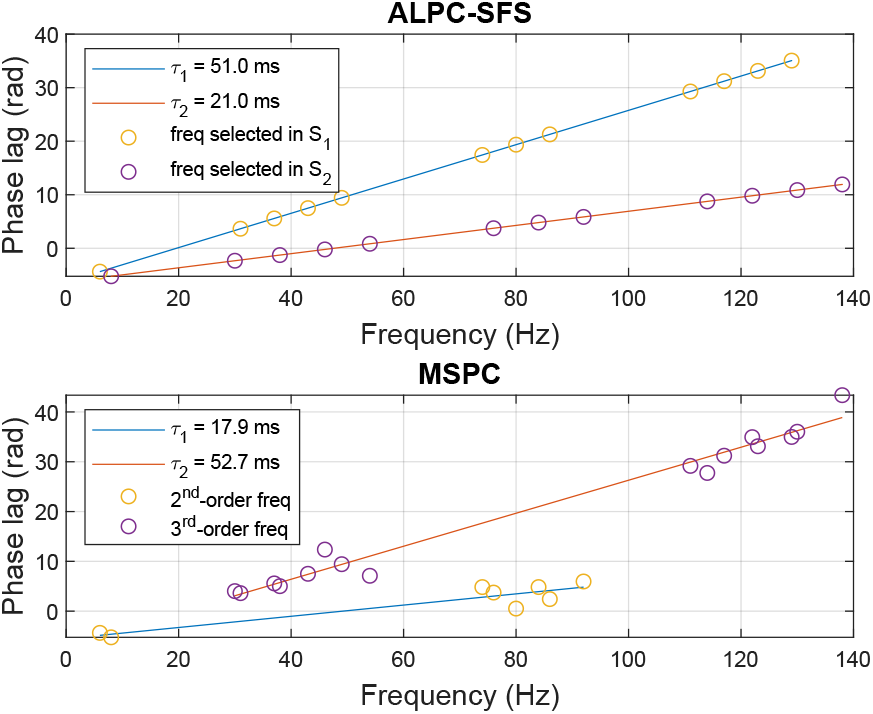
Estimated latencies when subsystems (*y*_1_ and *y*_2_) are unknown. (Top) By using ALPC-SFS on Y, two latencies (51 and 21 ms) are correctly estimated with corresponding MPEs less than 0.001 and with correctly identified subsystems (*S*_1_ = *y*_1_ and *S*_2_ = *y*_2_). Here, SFS started from a randomly selected distortion frequency, 6 Hz, and the same termination rule with scenario 1. (Bottom) MSPC estimated latencies for the assumed 2^*nd*^-order, and 3^*rd*^-order subsystems from X and Y. The estimated latencies are 17.9 ms and 52.7 ms with large MPEs of 0.78 and 0.66 for the 2^*nd*^-order and 3^*rd*^-order subsystems, respectively. Note also that the selected frequencies that determine the two latencies do not correspond to the actual subsystems.

#### 3) Example 3: two mixed subsystems with identical outputs

We also simulated the more challenging scenario in which the two subsystems had overlapping output components. In this case, only distorted phases can be extracted from the mixed output signals, and the relative amplitude gain of the output components (related to *ψ_mr_* in (1)) results to determine the estimated latencies. The stimulated model reads:

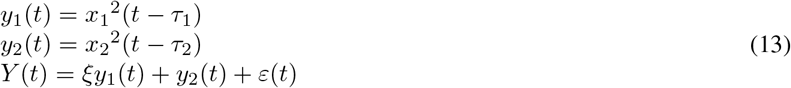

where the relative amplitude gain *ξ* > 0, and the latencies, *τ*_1_ = 15, *τ*_2_ = 20 ms. Both *y*_1_ and *y*_2_ had identical input frequencies (17, 21, 27) Hz so that they generated nine identical 2^*nd*^-order output components: {4, 6, 10, 34, 38, 42, 44, 48, 54} Hz. We varied the relative gain 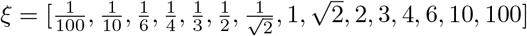, and simulated the mixed signals Y without noise, and under SNRs of (5, 0, −5, −10, −15, −20) dB. The latency estimates as function of the relative gain are shown in Fig. 8.

**Fig. 8.**
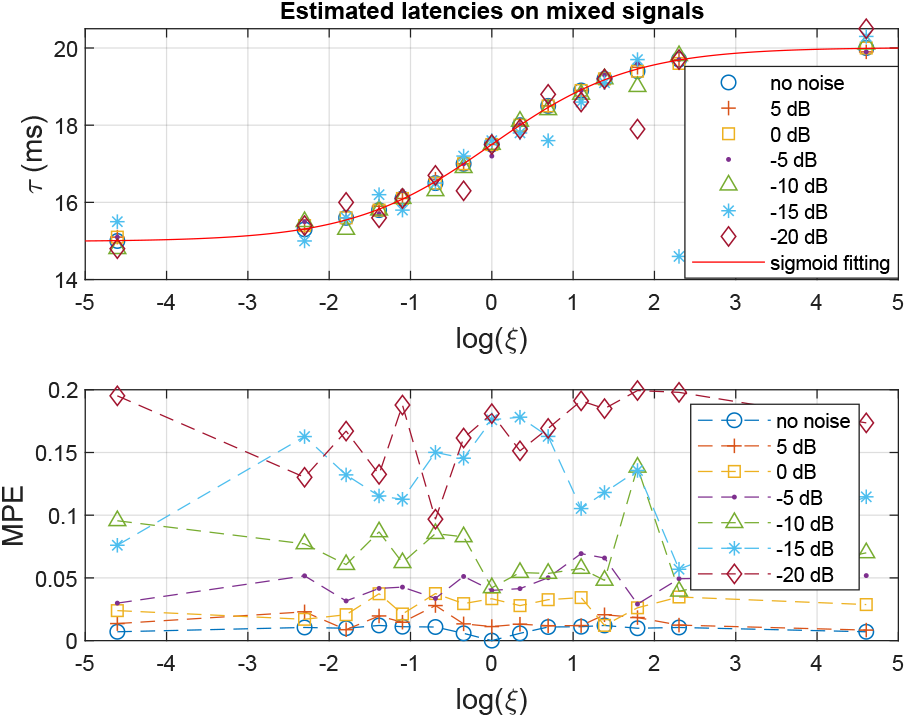
(Top) Estimated latencies on mixed signals of two subsystems with overlapping output frequencies and varying amplitude scale ratio *ξ*. The estimated latencies from mixed signals (on the noise-free signals) fit a sigmoid function between two veridical latencies. (Bottom) The mean phase error (MPE) tends to increase with decreasing SNR. In all cases, SFS started with 10 Hz.

The simulations show that in case of total overlap of the system outputs, the estimated latency is a weighted average of the underlying true latencies, where the relative gain of the systems serves as the weighting factor. After normalizing the estimated latencies (on the pure, noiseless signal) to a range between 0 and 1, we fitted the estimates with a sigmoid function 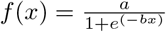, with x the natural logarithm of the relative gain, and fit coefficients a=1.00 and b=1.17, with 95% confidence bounds (0.9944, 1.007) and (1.143, 1.196), respectively. In general, the estimated latency for the superposition of the two system outputs with latencies (*τ*_1_ ≤ *τ*_2_) follows: 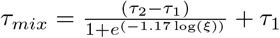, which is equivalent to,

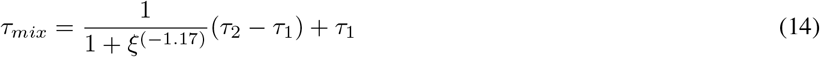

On the signal without noise, each selected starting frequency of the SFS yielded the same results for latency, MPE, and selected frequency components (i.e., output components). On the signal with noise, selecting a different starting frequency for the SFS yielded similar results for latency and MPE (as shown in Fig. 8), but could slightly differ regarding the selected frequency components, because frequency components with a large phase shift (due to low SNRs) are not selected by SFS.

### B. Measured ASSR components in the EEG

Table I shows the nonlinear interactions between the four stimulus input frequencies up to 6^*th*^ order, and the corresponding frequencies that could potentially be present in the measured ASSR. The odd-order distortion products (around 500 Hz and higher; e.g., 2*f*_1_ − *f*_2_) fall above the pre-processed EEG bandwidth (< 200 Hz). In this study, we used the same four frequencies (with different SPLs) as stimuli for both ears. As a result, all potential binaural beats (BBs, if significant) overlap with the monaural beats (MBs), whereas MBs often show much larger ASSR amplitudes than BBs ([40]). Therefore, the ASSRs analyzed in this study are the result from a superposition of MBs and potential BBs, but are mainly dominated by the MBs. This study therefore focused only on the method to estimate ASSR latencies, rather than to disentangle the contribution of MBs vs. BBs.

**TABLE I.**
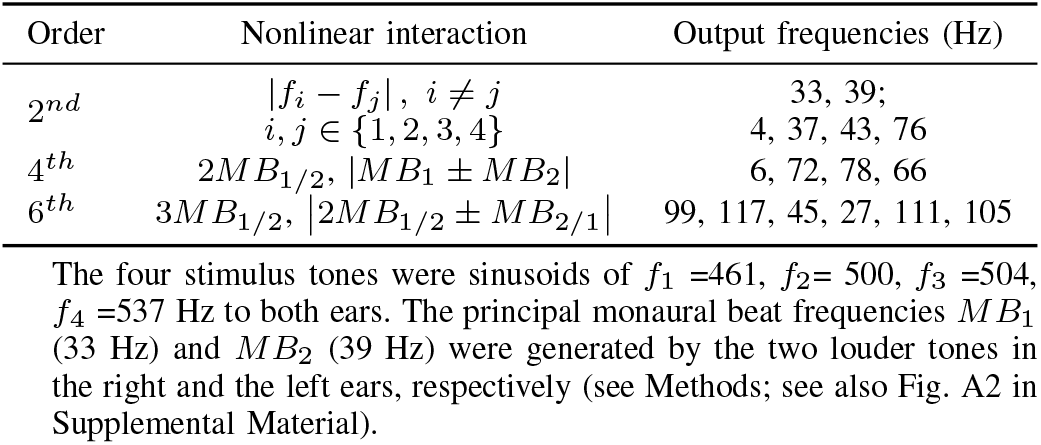
ASSR COMPONENTS GENERATED BY THE NONLINEAR INTERACTION

In the ASSRs, two predominant MBs (i.e., 33, 39 Hz) were generated by the second-order difference of the two louder pure tones in each ear. Their full 2^*nd*^-order interactions (including harmonics and their inter-modulations, i.e., a subset of 4^*th*^-order interactions of the input frequencies) and 3^*rd*^-order interactions (i.e., a subset of 6^*th*^-order interactions of the input frequencies) also show high SNRs. Note that Table I shows a subset (<200 Hz) of theoretical ASSR frequencies corresponding to each system order. In addition to the 4^*th*^-order output frequencies shown in Table I, more 4^*th*^-order output frequencies could potentially be generated among the full set of 2^*nd*^-order output frequencies, e.g., 29 and 35 Hz. Such output frequencies, however, were not considered in this study because of much lower SNRs.

Figure 9 shows that, in the measured EEG, the theoretical ASSRs all exceeded the background (i.e., non-ASSR integer frequency components) of 6.3 (±0.6) dB and the threshold of 7.492 dB (p=0.01). To avoid the potential detrimental effect of EMG artifacts, we averaged the top-3^*rd*^ SNRs on each integer frequency across ten subjects, measured on both recording days. The measurements of day 1 and day 2 showed highly similar SNR values. The grand average (n=20, day 1 and 2) SNR across all 61 EEG channels showed that ASSRs tended to have their maximum activation at both temporal lobes and frontal-central regions (bottom panel of Fig. 9). Results from the individual subjects are provided in the Suplemental Material, section D.

**Fig. 9.**
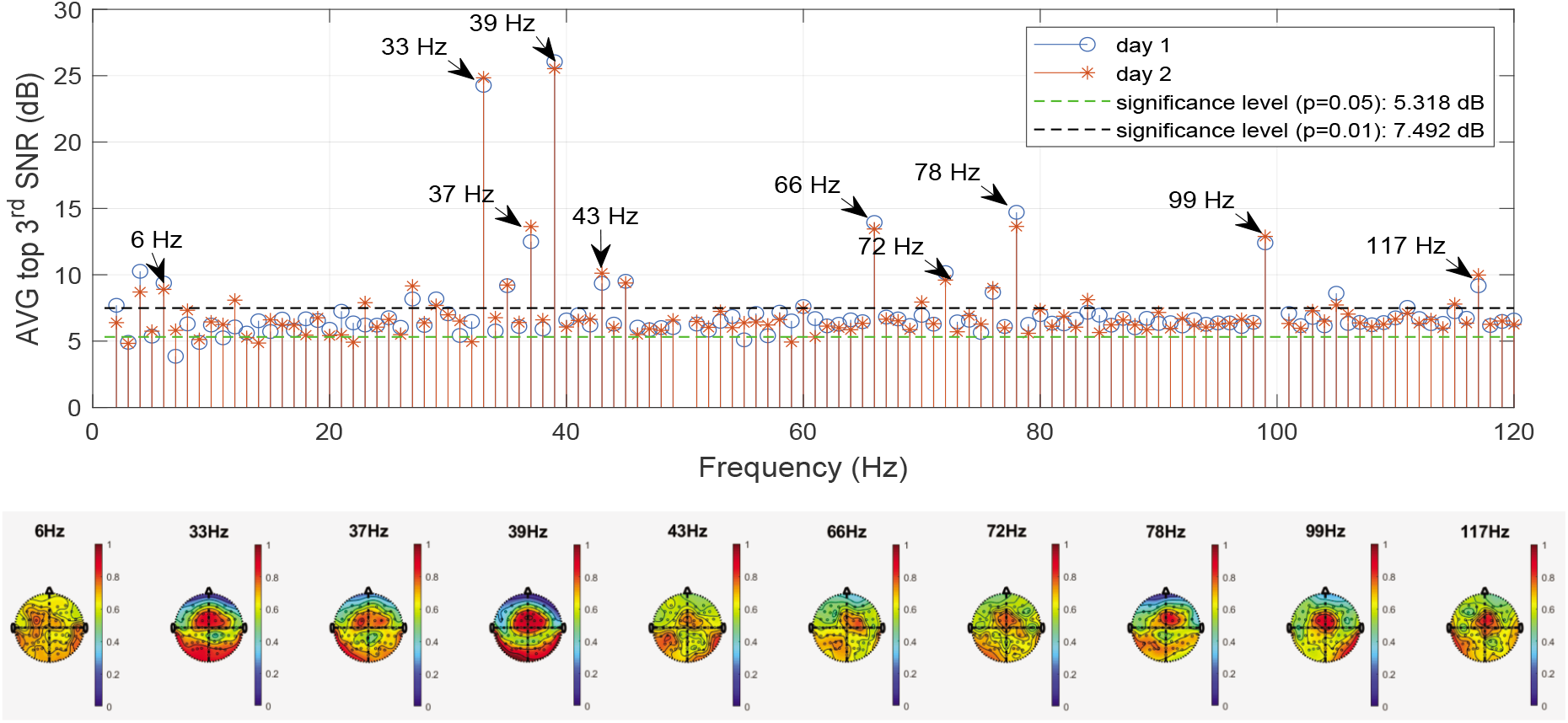
Grand average SNRs (n=10 subjects on recording days 1 and 2, top panel) and grand averaging scalp topography (n=20 measurements over two days, bottom panel) on ten major ASSR components indicated by arrows. The third-largest SNR (out of 61 channels) on each integer frequency was averaged across 10 subjects measured on day 1 or day 2. To calculate the grand average of the scalp topography, the SNR of each measurement was normalized between 0 and 1 to enable a direct comparison of each subject’s contribution.

### C. Estimated latencies from ASSRs

The MB frequencies showed a higher SNR (or LCI) than the other ASSR components. Therefore, in applying ALPC-SFS, we estimated latencies by starting with a pair of MB frequencies (i.e., 33 and 39 Hz). The MB frequency pair can determine a search direction for SFS, so that it finds the involved ASSR components that have a common latency. Similarly, we evaluated the underlying latencies dominated by higher frequencies (HF) (>80 Hz) by using SFS starting with the 3MB frequencies (i.e., 99 and 117 Hz). The ten subjects yielded similar results for the measurements on both days. The results of day 2 are shown in Fig. 10, where the phase extraction was performed by taking the ‘AVG EEG’ procedure (see Methods).

**Fig. 10.**
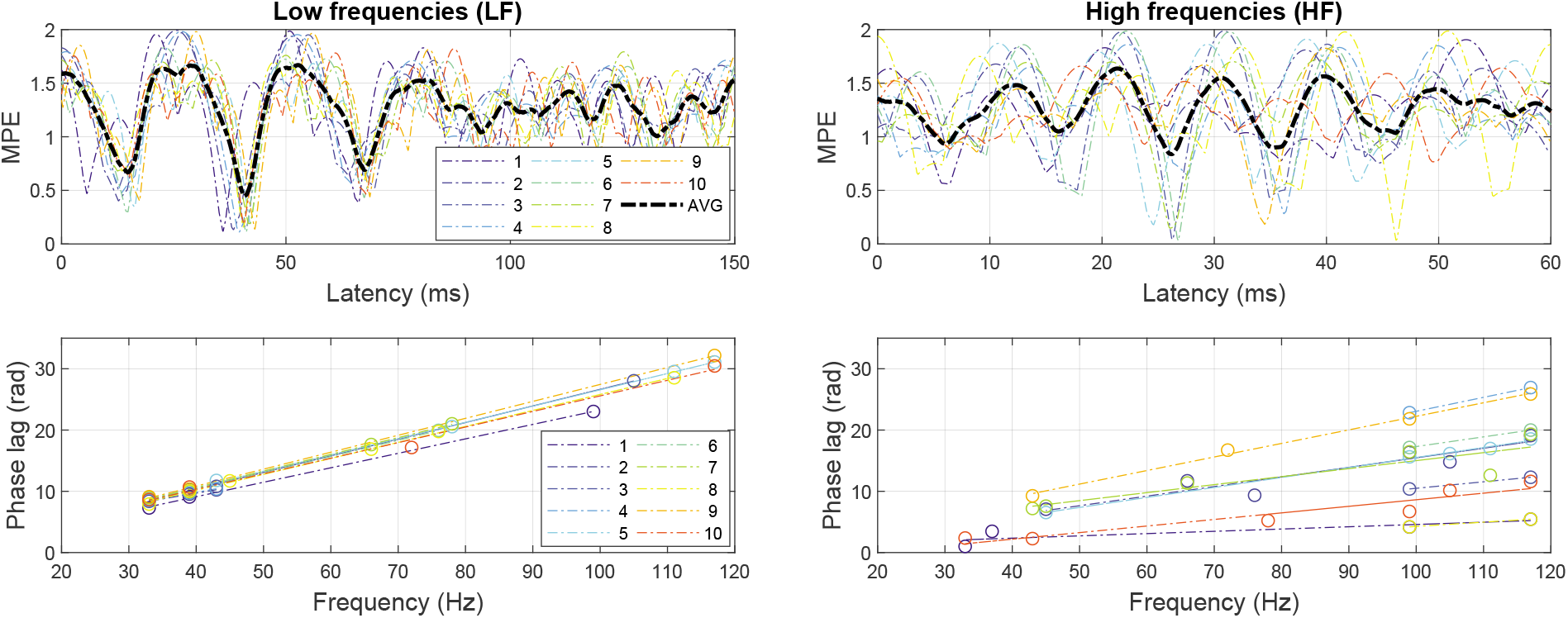
Estimated latencies of 10 subjects on EEG channel P8. The left-hand panel shows the latencies determined when SFS started with the MB frequencies (33 and 39 Hz), which overlap considerably because of a stable slope (latency estimate) across subjects; the right-hand panel shows the result when SFS started with the 3MB components. The minimum mean phase error (MPE) within the imposed prior range (<100 ms) corresponds to the estimated latency, which is proportional to the slope of the fitted lines in the bottom panels.

The estimated latencies resulted to vary with the location of the EEG channels. Table II includes the estimated latencies for three representative EEG channels: P7 (left TL), FCz (FC region) and P8 (right TL). Channels FCz and P8 were chosen because they yielded the largest grand-average LCI. P7 is located contralateral to P8, and was selected for comparison.

**TABLE II.** ESTIMATED LATENCIES, MPE, AND LCI ON LF- AND HF-DOMINATED SYSTEMS (N=20).

| EEG channel | LF (m, ±sd) | | | HF (m, ±sd) | | |
| --- | --- | --- | --- | --- | --- | --- |
|  | Latency* | MPE | LCI (MBs) | Latency* | MPE | LCI (3MBs) |
| P8 (right TL) | 40.8 (38.8, 41.3) | 0.14 (±0.03) | 0.29 (±0.12) | 26.7 (25.2, 34.6) | 0.11 (±0.06) | 0.056 (±0.032) |
| FCz (FC) | 51.7 (27.6, 54.4) | 0.16 (±0.05) | 0.37 (±0.08) | 20.6 (16.7, 26.8) | 0.14 (±0.04) | 0.076 (±0.044) |
| P7 (left TL) | 42.5 (40.6, 68.2) | 0.13 (±0.05) | 0.26 (±0.14) | - | - | 0.047 (±0.038) |

TL: temporal lobe; FC: frontal central. MPE: mean phase error at the estimated latency. $MPE \in [0, 2]$ . Note the low values for the MPE, indicating high confidence in the estimates.
\* Median values, and 25% and 75% percentiles, respectively.

The results based on AVG phase and AVG EEG were quite comparable (see Fig. 11). The estimated latencies for the low-frequency distortion products (LF) on EEG channels FCz and P7 had a non-Gaussian distribution, so that we report their median values and percentiles. The MPEs at the estimated latencies were low (<10%) and at a similar level for all estimates of the two LF- and HF-dominated systems, which is indicative for a reliable estimation. The most consistent latencies were obtained from EEG channel P8, which yielded a large ASSR response (i.e., high SNRs and LCIs). As the power of high-order responses (e.g., 4^*th*^ and 6^*th*^ orders) tended to decrease, the corresponding ASSR components were not significant on certain EEG channels, causing a less reliable estimated latency. We thus selected a subject-specific best channel with the largest LCI value instead of using P7 for estimating the HF-dominated latencies, showing a median latency of 21.1 ms, with 19.9 and 24.3 ms as the 25% and 75% percentiles, respectively. The ‘best ch’ is mainly selected from the FC region, thus it shows a similar median value as FCz. The HF-dominated latency generally showed a larger variance than the LF-dominated latency, which relates to a smaller LCI of the HF components.

**Fig. 11.**
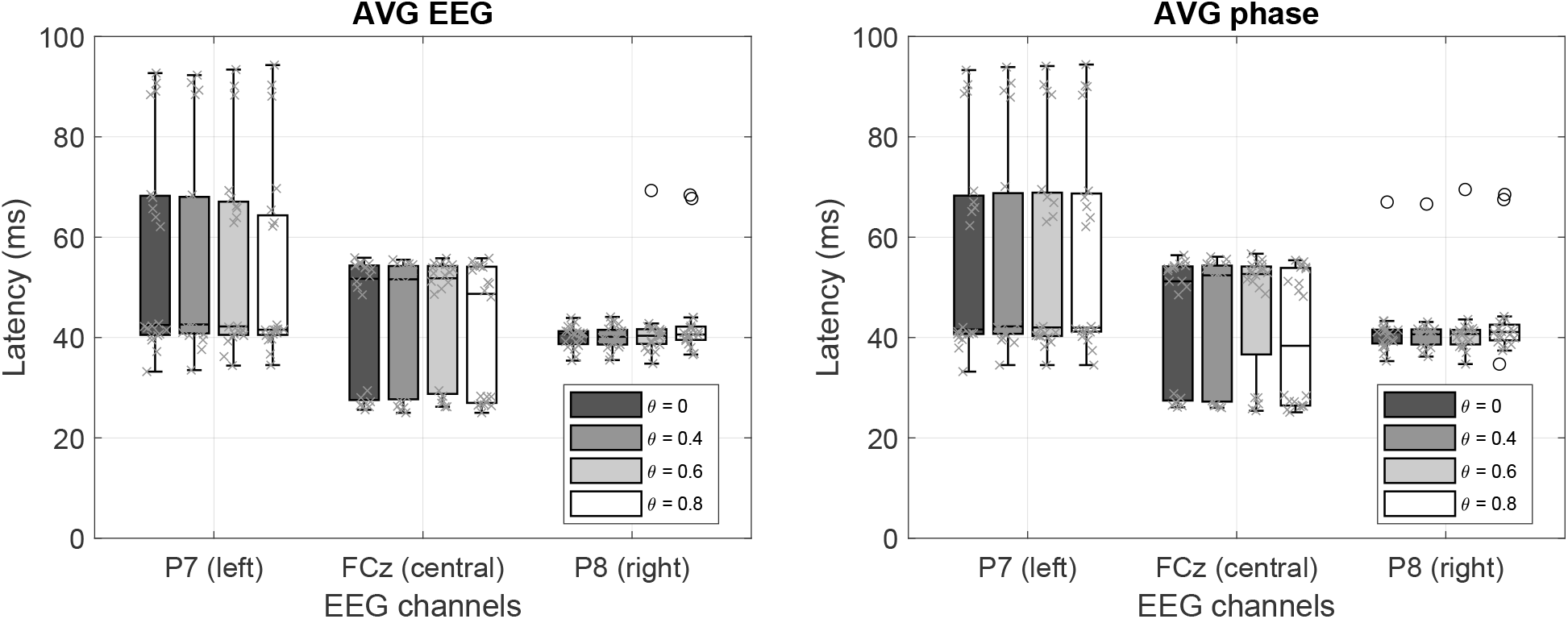
Effect of epoch filtering (LF-dominated latency as an example (n=20)). We applied EEG epoch (1 sec) filtering with four thresholds of the PLV (*θ* = 0, 0.4, 0.6, and 0.8). Both phase-extraction methods (AVG EEG and AVG phase) yielded highly similar latency estimates.

The LCI was computed on K=1200 (100 trials*12 sec) EEG epochs with 1-sec duration for each ASSR component. According to the theoretical threshold of 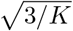 (see Methods), the 95% confidence threshold for a significant LCI was 0.05. We used the average LCI of MBs and 3MBs for the LF and HF components. Table II shows that the LCIs for LF components were significantly higher than for HF components, which is mainly due to the decreased SNRs of the higher-order ASSRs (see Fig. 9). The lower LCIs on the HF components partly explains the larger variance on the estimated latencies.

### D. Effect of epoch filtering

We excluded poorly phase-locked epochs by applying epoch filtering (see Methods) and compared the results for four different thresholds (*θ* = 0, 0.4, 0.6, and 0.8) for the phase-lock values (PLV). These PLV thresholds were chosen empirically, (*θ* = 0 means that 100% of the EEG epochs were included), such that the included EEG epochs were around 60%, 30% and 10% for *θ* = 0.4, 0.6, and 0.8, respectively. The estimated latencies were robust for different values of *θ*, as shown in Fig. 11. The cause for this stability is that the ALPC-SFS method employs multiple ASSR frequencies to determine a latency, and among them the MB frequencies show generally stable phase values across subjects. As a result, the Average EEG and Average Phase extraction methods both yielded similar results. Around 10% of all EEG epochs (i.e., *θ* = 0.8) generated similar results to that using all EEG epochs, which indicates that the latency is mainly determined by EEG epochs that are strongly phase-locked to the stimuli. On the other hand, using fewer EEG epochs increased the variance of estimated latency across subjects.

### E. A comparison with the MSPC method

We compared our ALPC-SFS method with the MSPC method for estimating the ASSR latencies from the recorded EEG. Figure 12 shows the results obtained from one representative subject. The ALPC-SFS method selects a group of ASSR frequencies that share a common latency across different potential system orders (panel b). In contrast, the MSPC method computes a latency by assuming a specific system order. All significant ASSRs (i.e., LCI > 0.05) were used. Note that the LCI is mathematically equivalent to the phase-coupling strength employed in the MSPC method (see Methods). The latter method, however, led to larger MPEs, because the ASSRs of the same order could have been generated by subsystems with different latencies, yielding poorer linear regression results for the relation between phase lag vs. frequency (see Fig. 12).

**Fig. 12.**
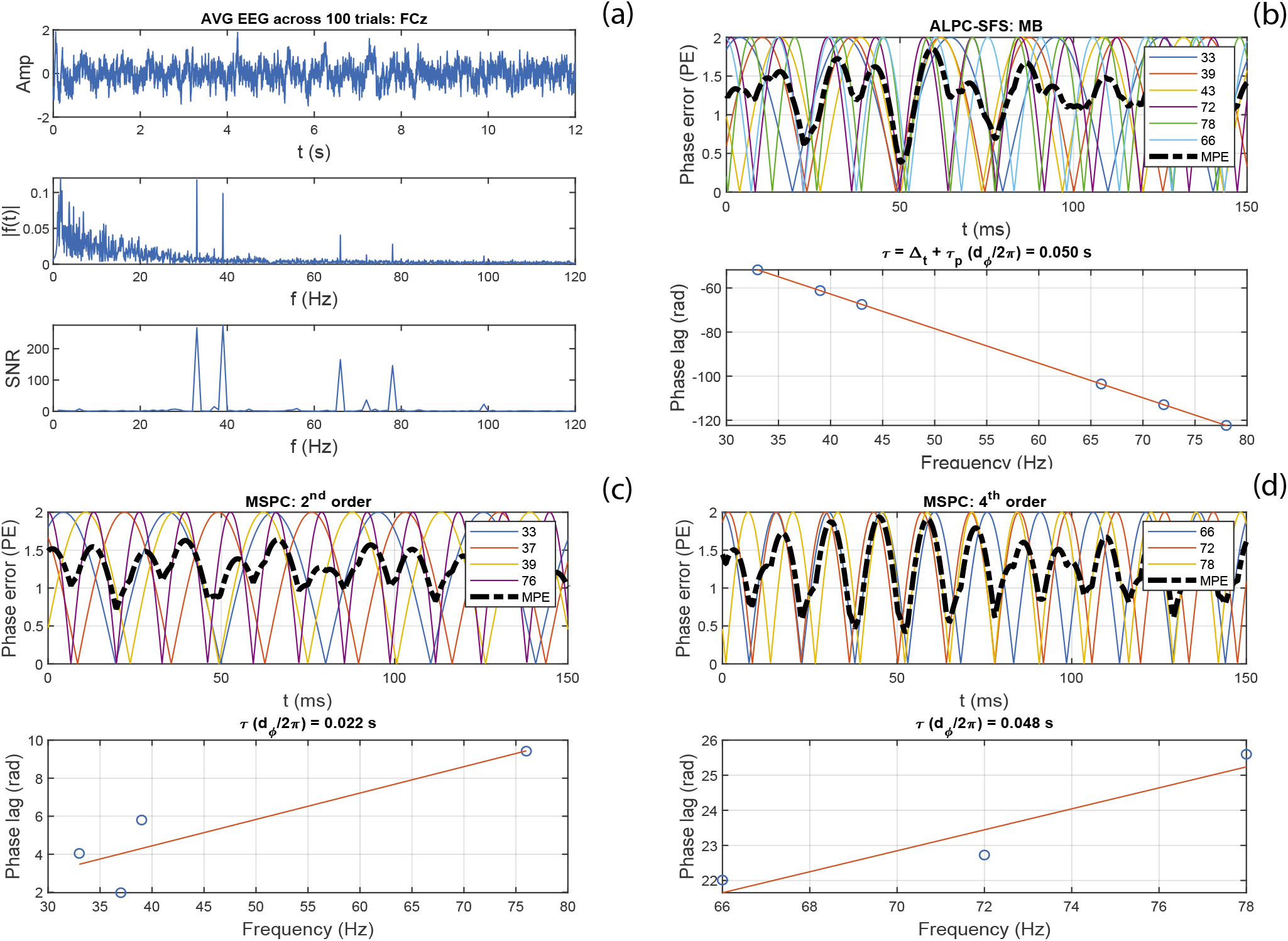
An example using one channel EEG signals (FCz, 100 trials) from Subj#1 to demonstrate the difference in results between the ALPC-SFS and MSPC methods. (a) shows the average EEG across 100 trials, its spectrum, and the neighboring SNR (see Methods). (b), ALPC-SFS starts from two MB frequencies (33, 39 Hz), and automatically selects the ASSR frequencies that have a similar latency (error < 5 ms), leading to a MPE < 0.5. ALPC-SFS applied time compensation, which leads to the negative slope that corresponds to the pseudo latency. The MSPC method estimated the latencies by assuming specific system orders (panel c: order 2) and (panel d: order 4) (see also Table I). However, the lumped 2^*nd*^-order ASSRs did not yield a reliable latency estimate, although all selected ASSRs were significant (i.e., LCI > 0.05). In contrast, the ALPC-SFS procedure identifies its latencies across system orders, and generated a subset of ASSRs with the same latency from both 2^*nd*^ order (33, 39, 43 Hz) and 4^*th*^ order system outputs (66, 72, 78 Hz).

Table III shows the estimated latencies from the MSPC method for 20 measurements on two EEG channels: FCz and P8, on which subjects showed generally more significant ASSRs. Compared with the ALPC-SFS based MPE values in Table II, the MPE values (> 0.3) in Table III indicate that the estimated latencies from the MSPC method were not reliable. In addition, the low LCI values show that many ASSRs were weak and not significant (i.e., LCI < 0.05), and were excluded for estimating latencies. Note that the median values of the ALPC-SFS vs. MSPC-based latencies were similar (e.g., when based on 2^*nd*^-order outputs). The ALPC method becomes mathematically equivalent to the MSPC method when the assumed system information is correct. However, compared with the ALPC-SFS method, the MSPC method lacks the procedure of ASSR frequency selection, and instead relies on a prior assumption regarding the system order to group frequencies (see Fig. 12). Such an assumption may be appropriate for the skeletal motor system ([20]), but may be less suitable for analyzing the human auditory system. For example, looking at the 4^*th*^-order distortion products, the MSPC method would include 2MBs, which are already generated at the cochlea, as well as the difference frequencies (e.g., 6 Hz) of MBs from both ears, which are generated at higher levels in the binaural auditory pathways (e.g., at the inferior colliculus, or at auditory cortical levels). Possibly, these different components may have different latencies, and should not be lumped into the same subsystem for estimating a latency.

**TABLE III.** Estimated latencies with the mspc method for three system orders

| System order | FCz |  |  | P8 |  |  |
| --- | --- | --- | --- | --- | --- | --- |
| | Latency* | MPE (m, ±sd) | LCI (m, ±sd) | Latency* | MPE (m, ±sd) | LCI (m, ±sd) |
| $2^{nd}$ | 55.3 (52.2, 84.7) | 0.68 (±0.24) | 0.16 (±0.02) | 40.7 (15.8, 41.5) | 0.63 (±0.25) | 0.12 (±0.04) |
| $4^{th}$ | 40.3 (37.1, 50.4) | 0.31 (±0.19) | 0.07 (±0.04) | 42.3 (32.2, 55.5) | 0.48 (±0.26) | 0.06 (±0.03) |
| $6^{th}$ | 22.0 (12.1, 31.1) | 0.38 (±0.22) | 0.05 (±0.02) | 45.2 (25.2, 46.6) | 0.33 (±0.25) | 0.04 (±0.02) |

\* Median value, and the 25% and 75% percentiles.
MPE: mean phase error, $\in [0, 2]$ , at the estimated latency.
LCI here is mathematically equivalent to the MSPC value used in [20].

## V. Discussion

### A. Advantages of the proposed method and stimuli

ALPC combined with SFS and time compensation has three major advantages compared with existing methods: (i) It requires no prior assumptions of the underlying nonlinear systems (e.g., system orders, or the number of subsystems) to calculate the unknown phase information of the ASSR components through phase compensation, and is therefore easier to implement. (ii) It allows for the estimation of multiple underlying latencies. (iii) It allows for the analysis of different nonlinear (sub)systems with different nonlinearities in one pass, and automatically selects the contributing ASSR components with the same latency (and potential common generators). Therefore, ALPC-SFS is better suited for analyzing nonlinear, higher-order ASSRs that are thought to underlie complex brain functions.

Moreover, the use of multi-cosine stimuli with all initial phases set to zero provides two additional advantages. (i) Compared to stimuli with arbitrary initial phases, it avoids the complicated procedure of phase compensation, and thus requires no prior assumptions regarding the underlying system nonlinearities. Note that the stimuli should preferably be cosine (not sine) with initial phase at zero, because for multi-cosine stimuli, arbitrary-order (even and odd) nonlinear distortion products all have zero initial phase, so that one can readily apply ALPC-SFS with time compensation. As shown in Appendix C of the Supplemental Material, a sum of sines only maintains a common initial phase for a specific subset of distortion products only, because the nonlinear distortions of the sine will generate phase shifts ∈ [0, *π*/2, *π*, or 3*π*/2] that depend on system order and nonlinear interactions. (ii) The estimated latency for zero-phase distortions will be more robust against the unavoidable small system errors (e.g., small delays caused by tubes and the measurement devices). It is often difficult to directly measure the exact phase values of the stimulus carriers at the onset of experimental trials, i.e., the exact time at which sound stimuli hit the ear drums. As a result, even a small timing error of one millisecond will already significantly change the phases of high-frequency stimuli (e.g., > 1k Hz). Consequently, small errors in the stimulus phases may lead to an unreliable latency estimate for methods that require exact initial phase values of the stimuli. In contrast, for multi-cosine stimuli the exact onset phase at the ears is not critical, as the estimated latency will be the sum of the true latency and the unknown (small) system delay, which can be readily incorporated as a fixed time compensation (described in Methods).

Therefore, to evaluate higher-order ASSRs, our proposed cosine stimuli with zero phase serve as a convenient alternative to multiple-sine stimuli with arbitrary initial phases; the ALPC-SFS method can also be applied to amplitude-modulated stimuli, provided that the initial phases of the modulation frequencies are all set to zero.

The ALPC-SFS method is a phase-based latency-estimation method. In addition to the phase-based methods, amplitude-based methods have been developed, in which the latency is estimated by the time shift that maximizes the statistical correlation between the envelope of the stimulus signal (e.g., a frequency modulation) and the response ([41]), ([42]). In general, phase-based latency-estimation methods are suitable when a limited number of ASSR frequencies is present. The amplitude-based methods are appropriate for broadband response signals, consisting of a continuous frequency spectrum. However, it is worth noting that in order to analyze higher-order ASSRs, phase-based methods may outperform amplitude-based methods, because the latter are applied to analyze the envelope of the stimuli and responses, which are limited to only 2nd-order nonlinear distortion products.

### B. Limitations of the method

The proposed ALPC-SFS method can estimate multiple ASSR latencies from a single EEG channel. However, it has two major limitations. First, the estimated latency is directly associated with ASSR frequencies, rather than underlying discrete neural sources. Thus, the method cannot disentangle the contributions from multiple neural generators. Instead, to reveal the potential ASSR sources from the estimated latencies, it will require prior information about the relationship between ASSR frequencies and potential neural sources. Second, when the same ASSR frequencies stem from multiple sources (which can be measured from the EEG), the ALPC-SFS estimate will fall between the two true latencies, where the exact value will be determined by the relative ASSR amplitudes of the original systems (Fig. 8).

One of our future goals will be to associate a potential latency change (even to infinite, i.e., no response) with (dys)function of the auditory system of the hearing impaired. To this end, in addition to estimate (and longitudinally follow) the ASSR latencies, we also need to evaluate the underlying neural generators. For that, it will be helpful to take advantage of the multiple electrode setup, and to employ sophisticated source-localization techniques and methodologies, which enable a better spatial resolution ([7]), Bayesian model averaging, ([43]), beamforming techniques ([44]), as well as multivariate source separation [45].

### C. Pitfalls of EEG montage

For the preprocessing of EEG signals, it is detrimental to re-reference all channels to an EEG channel that is not neutral to ASSRs, since a re-reference EEG channel containing strong ASSRs will affect all other EEG channels. For example, Fig. 13 shows phase values of all EEG channels at one MB frequency. After re-referencing to EEG channel Cz (carrying a large SNR of ASSRs), the distribution of SNR and phase values clearly changed. The phase of all EEG channels became similar to that on Cz. Therefore, it is recommended to either use the CAR montage ([34]), or to re-reference to a neutral electrode. On the other hand, also the CAR montage has its drawbacks, as it may cause phase ambiguity on the EEG channels on which ASSRs are relatively low. Certain ASSR components might be reversed, resulting in a phase shift of π. For that reason, the EEG channel P7 shows a larger variance on the estimated latency. To avoid the influence of such phase ambiguity, it is good practice to use the EEG channels that show relatively stronger ASSRs, e.g., at the frontal-central (FC) and the right temporal-lobe regions. Note that completely reversing an EEG channel (i.e., y = -x) will cause the same phase shift of *π* on all frequency components. It thus only changes the bias term of a fitted line (phase lag vs frequency) without changing the slope. Therefore, it does not affect the results of ALPC, or apparent latency.

**Fig. 13.**
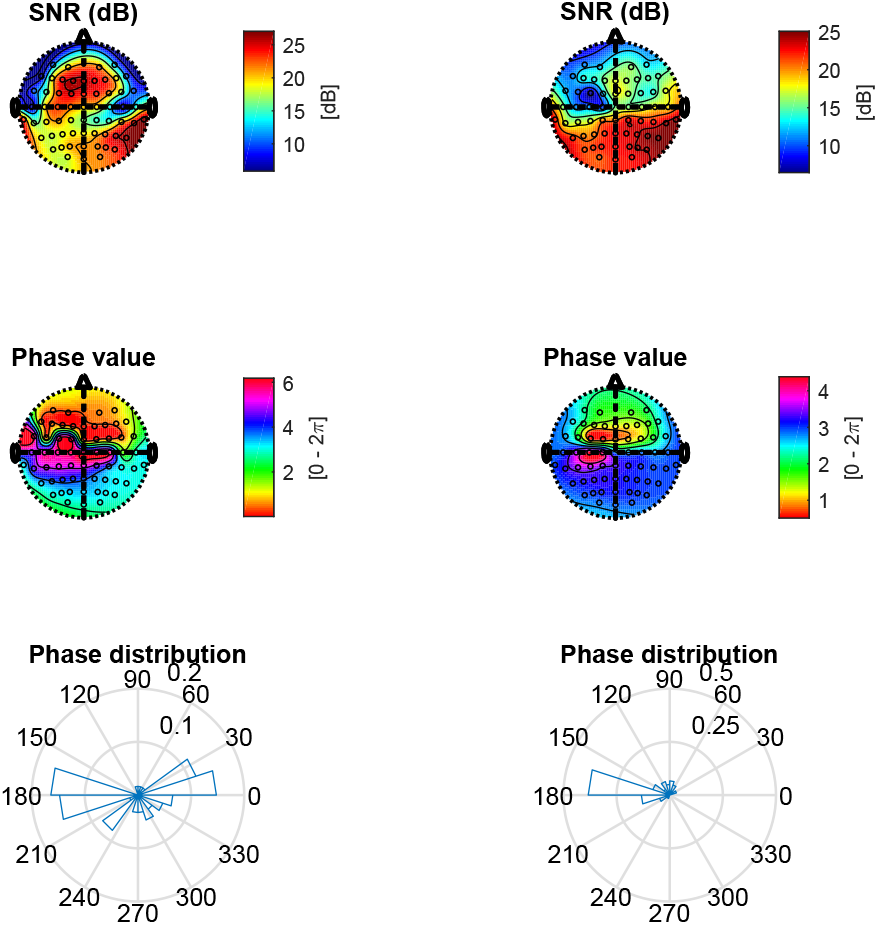
SNR and phase topography of the average EEG of 100-trials at an MB frequency of 33 Hz (data from subj#l). The left-hand column shows the common average reference (CAR) montage, while the right-hand column shows the result after re-referencing to EEG channel Cz. Note that the phase topography of the CAR shows a typical fronto-occipital phase delay, which is in line with ([46]). In contrast, the phases of many EEG channels (right-hand column) are changed by Cz, thus may leading to biased estimate of latencies.

### D. Multiple latencies of ASSRs

To estimate reliable latencies with the ALPC-SFS method, several important issues should be considered: (i) select the starting parameters for the SFS; (ii) use only EEG channels with a relatively high SNR (or LCI), and (iii) use EEG epochs that show high phase-locked values. An important parameter for the SFS is the starting frequency. SFS is a suboptimal method, and the starting frequency (or frequency pair) determines the direction of the heuristic search. It is good practice to start with those ASSR components with the highest SNR (or LCI), like the two major MB frequencies (33 and 39 Hz) in our experiments. Our simulation results demonstrate that the SNR of ASSRs affect the probability that two subsystems can indeed be separated (see Table AI). Starting the SFS with these two MB components also identified the other ASSR components with the same latency. We also started the SFS from 2MBs or from the (4, 6) Hz pair. However, only when the SFS started from the major MBs or from 3MBs, we obtained consistent results across subjects. ASSRs at 4 and 6 Hz are weak in most subjects, so that no reliable phase values are available for estimating the latency.

ASSR sources locate in multiple cortical as well as subcortical regions: ASSRs (< 20 Hz) originate from the auditory cortex; ASSRs around 40 Hz may arise from multiple locations including the brainstem, thalamus and the auditory cortex, while high frequency ASSRs (80 - 100 Hz) are supposed to dominate from the upper region of the brainstem [2], [47]. Our analysis revealed two major latencies from the ASSR components. The LF-dominant system (near 40 Hz) showed longer latencies than the HF-dominant system (>80 Hz). This finding corroborates previous studies using amplitude modulation (AM) stimuli on EEG or MEG signals [1], [22]. LF and HF -components correspond to the frequency ranges where maximum activation of ASSRs are thought to arise from the cortex (≈45 Hz) and the brainstem (≈ 90 Hz), respectively [27]. For the LF-dominant system, the two TL regions showed similar latencies (≈ 41 ms) whereas the FC region was endowed with a slightly longer latency (≈ 52 ms). This observation is consistent with the results from an EEG-based, binaural hearing study [46]. A longer latency suggests an origin in the upper stages of the ascending pathways of the auditory system. Thus, the longer latency in the FC region could perhaps arise from binaural interactions in the cortex, following monaural interactions. For the HF-dominant system, however, the estimated latency on the FC region (near FCz) was shorter than at both TL regions (P7 and P8), which is similar as reported by [22]). For these HF components, the brainstem has been considered as the major underlying source [27]. In the monaural auditory pathway, the signal may propagate from the lower brainstem, through the upper brainstem (which is anatomically located near the FC region), to the auditory cortex (located near the TLs).

It is further interesting to note that both LF and HF-dominant systems contained some overlapping ASSR components (see Fig. 10). This finding suggests that multiple neural mechanisms, characterized by different frequency ranges, may still enter a phase-locked status and thus shows a common latency. This finding is in line with the observed phase synchronization across multiple brain regions and function stages in the hearing pathways and high-level cortex [48], [3].

ASSRs are right-lateralized, i.e., right-hemisphere response exceeds the left-hemisphere response, which has been reported by several ASSR studies [7], [12]. We also found it in this study for both LF and HF ASSRs (see Table II), thus the EEG channel at P8 showed similar latency but smaller variance than on location P7 (see Fig. 11). In addition, the frontal-central (FC) brain region (near FCz) showed the strongest ASSRs. However, the latencies estimated from FCz had a larger variance than on P8. In particular, Fig. 11 shows two latency clusters, one major cluster around 52 ms and a smaller one around 25 ms. The major latency is longer than the TL region latency (around 40 ms), which might represent a binaural cortical interaction, i.e., the 4^*th*^ and 6^*th*^-order distortion frequencies are mainly generated from interaction between the beat frequencies of both ears. The smaller cluster (n=6 out of 20 measurements) should not be simply considered as outliers, because the FC brain region may represent the superposition of responses from multiple underlying sources (resulting in a weighted average latency estimate for brainstem and cortex). Furthermore, some subjects may have produced stronger ASSRs from the upper brainstem than from cortex, which would affect the estimated latency, as the brainstem-related latency is much smaller than the cortex-related latency [21], [49].

### E. Possible applications

The ascending auditory system, from auditory nerve to auditory cortex, contains several parallel pathways of acoustic signal processing, which eventually reach the cortical areas. For example, the localization of a sound source requires the neural processing of three types of acoustic cues, which are extracted by independent neural circuits within the auditory brainstem: interaural time differences (ITDs) and interaural level differences (ILDs) for locations in the horizontal plane, and direction-dependent spectral-shape cues from the pinna for directions in the vertical plane (Fig. 14; [50]).

**Fig. 14.**
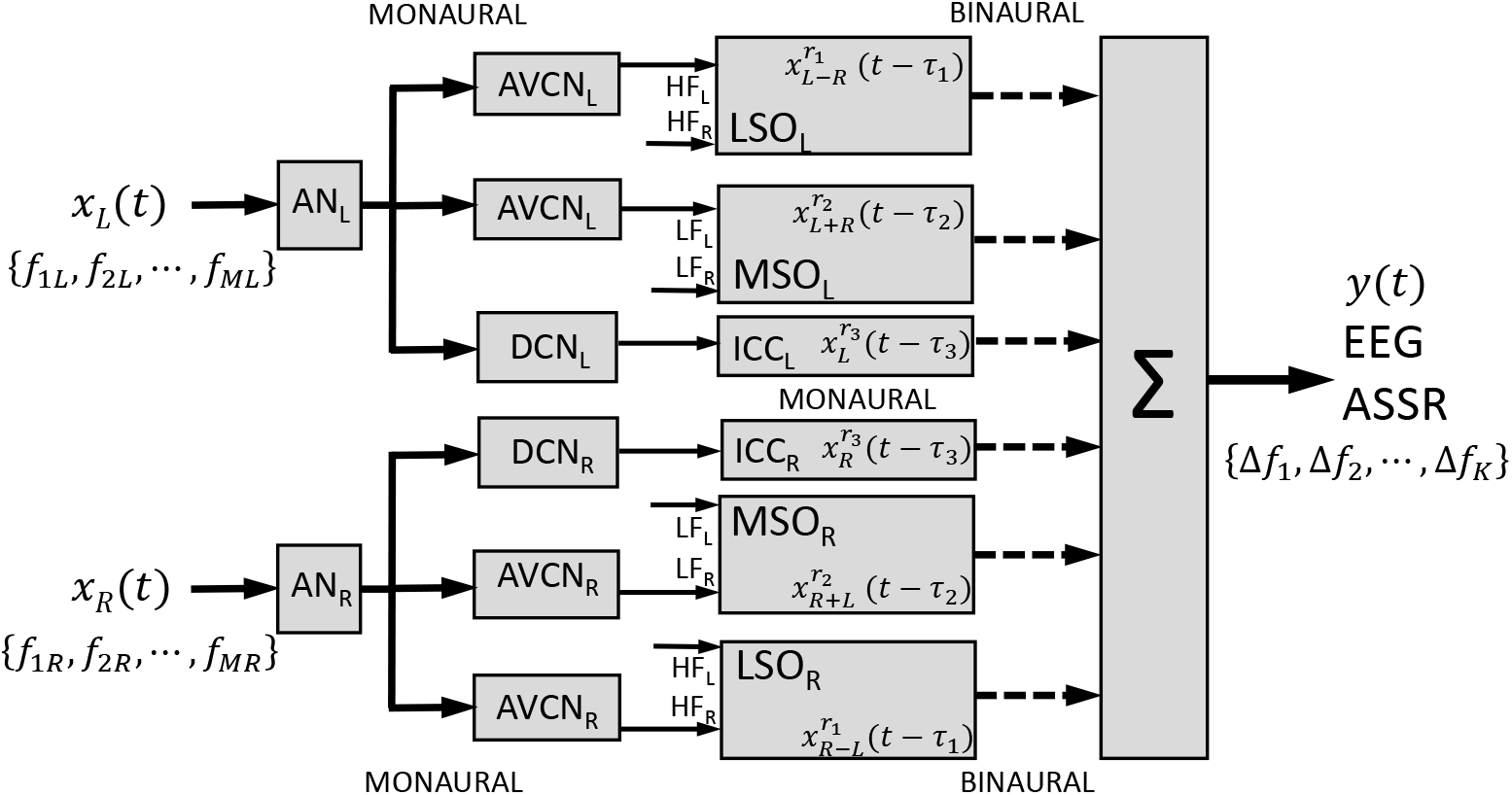
Sound localization relies on three independent monaural and binaural pathways that use largely non-overlapping frequency bands. All sounds are initially processed by a monaural stage at the auditory nerve (AN) and antero-ventral and dorsal subdivisions of the cochlear nucleus (AVCN and DCN). Low carrier frequencies (LF) are processed by the medial superior olive (MSO), which computes the interaural time difference (ITD); the high frequencies (HF) follow the lateral superior olive (LSO) pathway, which determines the interaural level difference (ILD). The DCN is part of a monaural path to the midbrain Inferior Colliculus (ICC) and cortex. All stages may introduce nonlinearities in the signal paths that are represented as unique frequency distortion products, Δ*f_n_*, in the EEG.

The binaural difference cues operate on complementary frequency bands: as the ITD for humans varies from about −600 μs (sound at the far left) to +600 *μs* (sound at the far right), this cue only provides unambiguous phase information for sound frequencies up to ~ 1500 Hz. For these low frequencies, phase-locked neurons in the auditory nerve, the anteroventral cochlear nucleus (AVCN), and the medial superior olive (MSO) carry precise phase information about the carrier frequency. In the MSO, neurons respond to the interaural phase difference at their characteristic frequency. Because the phase difference is frequency-dependent for a given delay (Δ*φ* = 2*πf* Δ*τ*), MSO neurons that are tuned to a specific frequency and phase difference thus encode a particular inter-aural absolute delay, and hence a particular direction in the horizontal plane. Recently, a periodic representation of ITDs has also been demonstrated for human cortex ([51]). Neurons that are tuned to the same direction in azimuth, but are sensitive to other frequencies, will be tuned to different phase differences. In this way, the neuronal processing resembles the ALPC algorithm, which also relies on frequency-dependent phase-differences to estimate the unique underlying (characteristic) delay ([52], [53], [50]).

A second pathway processes the ILDs that arise for frequencies above ~3 kHz because of the head shadow effect. Cells in the lateral superior olive (LSO) are tuned to the ILD for a particular frequency. The ILD cue depends strongly on the sounds frequency: the higher the frequency, the stronger the head-shadow effect. Combining the responses from a population of neurons, tuned to different frequencies, is needed to estimate the veridical azimuth angle of the sound source. It is conceivable that the final stage in estimating the sound’s azimuth angle involves nonlinear integrative mechanisms for both binaural pathways. Finally, the dorsal cochlear nucleus (DCN) feeds a monaural signal pathway via the central nucleus of the Inferior Colliculus (ICC) to the cortex, and is possibly involved in the localization of source elevation (spectral cue extraction), and pitch perception.

It is conceivable that the different auditory processing mechanisms may be endowed with different delays and different nonlinear processes. As the sound-localization mechanisms operate in different frequency bands, they could potentially introduce their own nonlinear distortions, and these may, in principle, be separated by our ALPC-SFS method, by applying specific frequency combinations to either ear. Similarly, the ALPC-SFS method may be used to test quantitative models of the ascending auditory system ([54], [19]).

In addition to evaluating the steady-state dynamics of the ASSRs (as done in the present study), the ALPC-SFS method could be extended to determine the latencies associated with the instantaneous phases of the ASSR frequencies, obtained by either Gabor wavelets or the Hilbert transform ([55], [56]). Such an extension could help to assess the neural mechanisms within the transition phases of the auditory (or visual) processing, as may be observed immediately after stimulus onset ([33]) or offset ([57]).

## VI. Conclusions

We proposed a nonparametric method, ALPC-SFS with time compensation, to estimate multiple latencies of the auditory system from ASSRs recorded on a single EEG channel. Compared with existing methods, our method requires no prior assumptions of the underlying nonlinearity (e.g., system orders), and can be readily implemented to analyze higher-order ASSRs, when combined with a zero-phase cosine stimulus complex. We illustrated our method on recorded EEG signals and it successfully identified two major (LF and HF-dominated) latencies that were stable across subjects and between measurement days. The LF-dominated latencies were longer than the HF-dominated latencies and may be putatively related to different brain regions (e.g., central brain or temporal-lobe regions). The LF-dominated latency is mainly determined by EEG epochs that are strongly phase-locked to the stimuli. ALPC-SFS is promising as an objective measure for the ASSR latencies and for the underlying nonlinearities of both primary and non-primary auditory cortex (dys)function.

## Acknowledgments

We thank Yuan Yang from Northwestern university for his advice on the experiments and for providing the analysis code of the MSPC method. We thank anonymous reviewers for their constructive criticisms on an earlier version of this manuscript, which greatly helped to improve our paper. Also, we thank the students and colleagues in Radboud university and Delft university for the help on collecting the EEG datasets. This research was supported by the Dutch Organisation for Scientific Research, NWO-TTW Perspectief, project ‘NeuroCIMT’ - Otocontrol, nr. 14689 (LW, EN), and EU Horizon 2020 ERC Advanced Grant 2016 ‘Orient’, nr. 693400 (AJVO).

